# Opioidergic activation of descending pain inhibitory system underlies placebo analgesia

**DOI:** 10.1101/2023.06.26.546410

**Authors:** Hiroyuki Neyama, Yuping Wu, Yuka Nakaya, Shigeki Kato, Tomoko Shimizu, Tsuyoshi Tahara, Mika Shigeta, Michiko Inoue, Kazunari Miyamichi, Natsuki Matsushita, Tomoji Mashimo, Yoshiki Miyasaka, Yasuyoshi Watanabe, Masayuki Kobayashi, Kazuto Kobayashi, Yilong Cui

**Affiliations:** Laboratory for Biofunction Dynamics Imaging, RIKEN Center for Biosystems Dynamics Research, Kobe, Japan; Department of Pharmacology, Nihon University School of Dentistry, Tokyo, Japan; Department of Molecular Genetics, Fukushima Medical University Institute of Biomedical Sciences, Fukushima, Japan; Laboratory for Comparative Connections, RIKEN Center for Biosystems Dynamics Research, Kobe, Japan; Laboratory for Pathophysiological and Health Science, RIKEN Center for Biosystems Dynamics Research, Kobe, Japan; Division of Laboratory Animal Research, Aichi Medical University School of Medicine, Aichi Japan; Division of Animal Genetics, Laboratiry Animal Research Center, Institute of Medical Science, The Universtiry of Tokyo, Tokyo, Japan; Laboratory of Reproductive Engineering, Institute of Experimental Animal Sciences, Osaka University Medical School, Suita, Japan

## Abstract

Placebo analgesia is caused by inactive treatment, implicating endogenous brain function involvement. However, the underlying neurobiological mechanisms remain unclear. We found that μ-opioid signals in the medial prefrontal cortex (mPFC) activate the descending pain inhibitory system to initiate placebo analgesia in neuropathic pain rats. Chemogenetic manipulation demonstrated that specific activation of μ-opioid receptor-positive (MOR^+^) neurons in the mPFC or suppression of the mPFC-ventrolateral periaqueductal gray (vlPAG) circuit inhibited placebo analgesia in rats. MOR^+^ neurons in the mPFC are monosynaptically connected and directly inhibit L5 pyramidal neurons that project to the vlPAG via GABA_A_ receptors. Thus, intrinsic opioid signaling in the mPFC disinhibits excitatory outflow to the vlPAG by suppressing MOR^+^ neurons, leading to descending pain inhibitory system activation that initiates placebo analgesia.

**One Sentence Summary:** Sugar pills relieve pain by activating the intrinsic pain inhibitory system via opioidergic signals in the prefrontal cortex.

## Main Text

Placebo effects are beneficial outcomes caused by ineffective sham medical treatment. Placebo effects exist in any clinical setting and affect therapeutic outcomes (*1*). However, their clinical applications have been limited because the underlying neurobiological mechanism remains unclear (*2*). Placebo analgesia is an actively studied placebo effect and a prime illustration of how the mind influences brain functions. The underlying mechanism of placebo analgesia involves activating the intrinsic descending pain inhibitory system by higher-order psychological processes (*3*). Neuroimaging studies in humans have demonstrated that hierarchical brain regions and neurochemical systems, such as dopaminergic and opioidergic systems, are involved in placebo analgesia (*4, 5*). Endogenous opioidergic systems at multiple brain levels have been implicated in placebo analgesia, including the rostral anterior cingulate cortex, amygdala, nucleus accumbens, and brainstem (*5-8*). However, the fundamental neurobiological mechanisms underlying placebo analgesia remain unclear owing to the lack of comparable neuroimaging results from animal studies that allow mechanistic exploration using molecular, cellular, and genetic manipulation.

Placebo analgesia is caused by expectancy and conditioning in humans and rodents (*9-12*). We have successfully identified several brain regions of rats, similar to those of human neuroimaging studies, which were involved in pharmacological conditioning-induced placebo analgesia using a small animal neuroimaging analysis and demonstrated that the endogenous μ-opioid system in the medial prefrontal cortex (mPFC) causally contributed to conditioning-induced placebo analgesia (*13*). The μ-opioid receptor (MOR) is expressed in GABAergic interneurons in the mPFC, and regulates the excitatory outflow of the PFC (*14*). However, the mPFC is connected to various brain regions and is engaged in diverse, complicated functions (*15, 16*); thus, whether and how the μ-opioid signaling in the mPFC controls the intrinsic pain modulation system for conditioning-induced placebo analgesia is unknown.

### Generation of MOR-Cre knock-in (KI) rat

To functionally manipulate MOR-positive (MOR^+^) neuron activity *in vivo*, we designed and developed genetically engineered rats expressing Cre recombinase under the control of the MOR 1 (*Oprm1)* promoter using CRISPR/Cas9 technology (fig. S1A). Double RNAscope^®^ *in situ* hybridization with probes targeting Cre and MOR gene showed that Cre signals were observed throughout the brain regions consistent with the MOR distribution (fig. S2), as previously reported based on ligand binding assay, and were almost wholly colocalized with the MOR gene not only in habenula and intrapeduncular nucleus but also in the mPFC (fig. S3; 98.9 ± 0.4%) (*17-19*).

### MOR^+^ neurons in the mPFC modulate neuropathic pain

To clarify whether and how MOR^+^ neurons in the mPFC modulate pain, we locally manipulated MOR^+^ neuron activity in the mPFC region of MOR-Cre KI rats with neuropathic pain using chemogenetic method and evaluated the pain behavior in response to nociceptive stimulation and related MOR^+^ neuron activity. We microinjected adeno-associated virus (AAV)-Flex-hM4Di-TagRFP or AAV-Flex-hM3Dq-TagRFP into layer 5 (L5) of the right prelimbic area of MOR-Cre KI rats, which is the major region of mPFC excitatory outflow (Fig. 1, A and I) (*20*). Three weeks later, TagRFP-positive (TagRFP^+^) neurons were observed around the L5 mPFC and in the L2/3, indicating Cre-dependent recombination of the transgene in MOR-Cre KI rats (Fig. 1C and J). As shown in Fig. 1D, most TagRFP^+^ neurons around L5 mPFC contained vesicular GABA transporter (vGAT) (73.1 ± 1.3%, 156 ± 14.0 cells from three rats), indicating MOR is largely expressed in the GABAergic interneurons, at least in L5 mPFC, as previously reported (*21*). Cre and TagRFP^+^ neurons were not detected in wild-type rats (fig. S4).

**Fig. 1.**
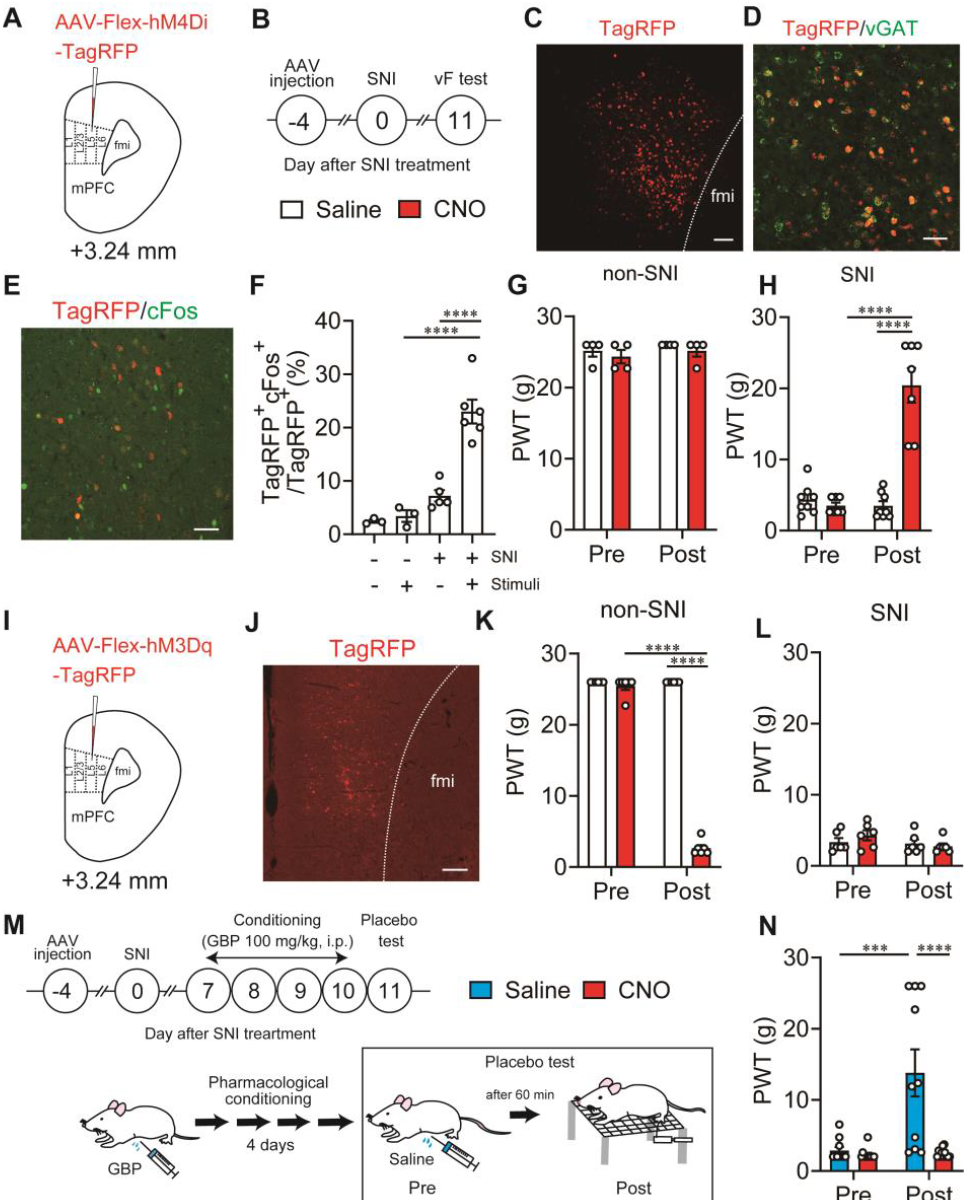
MOR^+^ neuron in the mPFC modulates placebo analgesia. (**A and I**) Schematic diagram of chemogenetic manipulation of MOR^+^ neurons of the mPFC of MOR-Cre KI rat. (**B**) Experimental procedure. (**C and D**) Representative mPFC coronal sections labeled with TagRFP (=MOR neurons, red) and vGAT (green) ISH probes. Scale bar: (**C**) 100 μm (**D**) 50 μm. (**E**) Immunostaining image showing cFos (green) expression in MOR^+^ neurons (TagRFP, red) in the mPFC. Scale bar: 50 μm. (**F**) The ratio of cFos^+^ neurons among MOR^+^ neurons. (**G**) Changes of paw withdrawal threshold (PWT) after saline and CNO injection in non-SNI rats microinjected with AAV-Flex-hM4Di-TagRFP. (**H**) Changes in PWT after injection of saline and CNO in SNI rats microinjected with AAV-Flex-hM4Di-TagRFP. (**J**) Immunostaining image of MOR^+^ neurons (TagRFP, red). Scale bar: 200 μm. (**K**) Changes in PWT after saline and CNO injection in non-SNI rats microinjected with AAV-Flex-hM3Dq-TagRFP, (**L**) Changes in PWT after saline and CNO injection in SNI rats microinjected with AAV-Flex-hM3Dq-TagRFP. (**M**) Experimental procedures for pharmacological conditioning-induced placebo analgesia and chemogenetic manipulation. (**N**) Changes in PWT after saline and CNO injection for examining placebo analgesia. \*\*\**P*<0.001, \*\*\*\**P*<0.0001. All statistical information details were presented in supplementary table.

To induce chronic neuropathic pain in MOR-Cre KI rats, we induced spared nerve injury (SNI) on the left hindlimb, which has been widely used as a neuropathic pain animal model (*22*) with persistent and stable pain hypersensitivity for at least 2 weeks (fig. S5, A and B). In these SNI rats, cFos and TagRFP double-positive neurons were significantly increased in the right mPFC in response to pain stimuli to the left hind paw 11 days after SNI treatment but not after sham treatment. These results indicate that MOR^+^ neurons in the mPFC were activated by pain stimuli (Fig. 1, E and F). Meanwhile, the chemogenetic inhibition of MOR^+^ neuron in the mPFC significantly suppressed the pain hypersensitivity (increased paw withdrawal threshold (PWT)) after intraperitoneal (i.p.) injection (1 mg/kg) of clozapine-*N*-oxide (CNO) in SNI rats expressing AAV-Flex-hM4Di-TagRFP in MOR^+^ neurons in the mPFC, but not in non-SNI rats (Fig. 1G and H). In contrast, the chemogenetic activation of MOR^+^ neurons by CNO (1 mg/kg) enhanced the pain responses in non-SNI rats expressing AAV-Flex-hM3Dq-TagRFP in MOR^+^ neurons in the mPFC, but not in SNI rats (Fig. 1, K and L). The chemogenetic activation of AAV-Flex-hM3Dq-TagRFP expressing MOR^+^ neurons in the mPFC was confirmed by cFos immunohistochemistry (fig. S6, A to C) and unit recording (fig. S6, D and E). Overall, MOR^+^ neurons in the mPFC are activated in response to pain stimuli and specific inhibition of this activity can produce an analgesic effect.

### MOR^+^ neurons in the mPFC modulate placebo analgesia

Subsequently, we examined whether and how MOR^+^ neurons in the mPFC modulate placebo analgesia. Placebo analgesia was induced by pharmacological conditioning in SNI rats, in which administration of pain-killer gabapentin (unconditioned stimulus) was paired with the i.p. injection (conditioned stimulus) for four consecutive days (Fig. 1M), as previously reported (*13*). On the test day (day 5), pain responses were significantly reduced by i.p. injection of saline as a placebo, indicating conditioning-induced placebo analgesia in these rats (Fig 1N, blue bar, increased PWT). In contrast, the chemgenetic activation of AAV-Flex-hM3Dq-TagRFP expressing MOR^+^ neurons in the mPFC significantly inhibited placebo analgesia after i.p. injection of CNO instead of saline (Fig 1N, red bar), but not in rats expressing AAV-Flex-TagRFP (control vector) (fig. S7). Overall, activating MOR^+^ neurons in the mPFC disturbs placebo analgesia.

### mPFC-ventrolateral periaqueductal gray matter (vlPAG) circuit modulates placebo analgesia

The mPFC sends monosynaptic excitatory projections into the ventrolateral periaqueductal gray (vlPAG), a key node nucleus of the descending pain inhibitory system (*23*), for modulating pain processing (*24-26*). Thus, we verified the functional role of the mPFC-vlPAG circuit in placebo analgesia. First, we confirmed the anatomical projections from the mPFC to the vlPAG. An anterograde tracing experiment showed that abundant enhanced green fluorescent protein (EGFP)-expressing fibers were observed in the vlPAG four weeks after microinjection of AAV-CaMKIIa-EGFP into the L5 mPFC to express EGFP in projection pyramidal neurons (PN) under the control of the glutamatergic specific calmodulin kinase (CaMKIIa) promoter (Fig. 2, A to D). Furthermore, retrogradely traced cholera toxin subunit B 647 (CTB647), was predominantly observed in L5PN of the mPFC after microinjection of CTB647 into the vlPAG region (Fig. 2, E to H). These results indicated that L5PN in the mPFC sends monosynaptic excitatory projections to the vlPAG region (*25*). That L5PN rarely overlapped with the MOR^+^ neurons (1.2 ± 0.3%) (fig. S8).

**Fig. 2.**
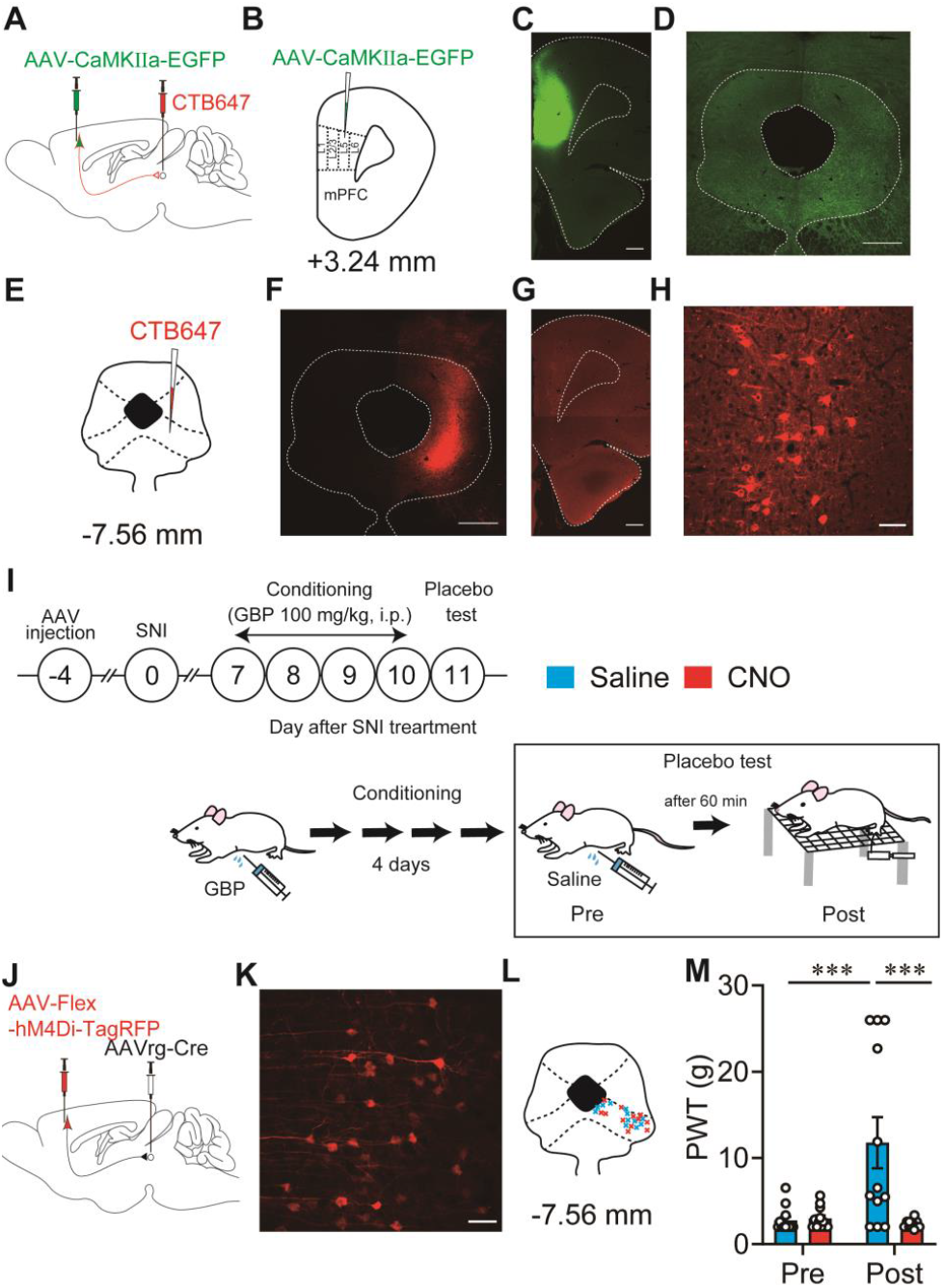
mPFC-vlPAG circuit modulates placebo analgesia. (**A and B**) Schematic diagram for anterograde (AAV-CaMKIIα-EGFP) and retrograde (CTB647) labeling of the mPFC-vlPAG circuit in wild type rat. (**C and D**) Images show AAV injection side in the mPFC and the projection fiber in the vlPAG that was anterogradely labeled. Scale bar: (**C**) 500 μm (**D**) 500 μm (**E to H**) Retrograde somatic labeling of L5 pyramidal neurons (L5PN) in the mPFC, in which CTB647 was injected into the vlPAG. Scale bar: (**F**) 500 μm (**G**) 500 μm (**H**) 50 μm. (**I**) Experimental procedures for pharmacological conditioning-induced placebo analgesia and chemogenetic manipulation. (**J**) Schematic diagram for chemogenetic manipulation of mPFC-vlPAG circuit using AAV-Flex-hM4Di-TagRFP. (**K**) High magnified image shows the expression of AAV-Flex-hM4Di-TagRFP in L5PN that projected into the vlPAG, blue, DAPI; red, TagRFP. Scale bar: 50 μm. (**L**) Retrograde AAV injection points in the vlPAG. (**M**) Changes in PWT after saline and CNO injection for evaluating placebo analgesia. \*\*\**P*<0.001. All statistical information details were presented in supplementary table.

Subsequently, we examined the functional role of that in neuropathic pain modulation. To specifically manipulate the mPFC-vlPAG circuit activity, we concomitantly microinjected retrograde AAV (AAVrg)-Cre into the vlPAG and AAV-Flex-hM4Di-TagRFP or AAV-DIO-hM3Dq-mCherry into the L5 of the right prelimbic area of the mPFC in wild-type rats (Fig. 2, I and J, fig. S9, A and E). Thus, AAV-DIO-hM3Dq-mCherry or AAV-Flex-hM4Di-TagRFP was specifically expressed in L5PN that sends monosynaptic projections from the mPFC to the vlPAG. (Fig 2, K and L; fig. S9, B and F). The chemogenetic activation of the mPFC-vlPAG circuit by CNO injection in rats expressing AAV-DIO-hM3Dq-mCherry significantly suppressed pain hypersensitivity in SNI rats (fig. S9D). However, it did not produce an analgesic effect in saline-injected control rats (fig. S9C). In contrast, the chemogenetic suppression of the mPFC-vlPAG circuit by CNO injection in rats expressing AAV-Flex-hM4Di-TagRFP significantly enhanced pain hypersensitivity in non-SNI rats but not in SNI rats (fig. S9G and H). Consistent with previous reports (*25-27*), these results indicated that the mPFC-vlPAG circuit modulates physiological and pathophysiological pain processing via the innervation of the downstream descending pain inhibitory system.

Finally, we confirmed whether the mPFC-vlPAG projection modulates conditioning-induced placebo analgesia. After four consecutive days of conditioning, the chemogenetic inhibition of the mPFC-vlPAG circuit specifically expressing AAV-Flex-hM4Di-TagRFP significantly reduced the analgesic effect (deceased PWT) in rats i.p.injected CNO instead of saline (Fig. 2M). The analgesic effect did not change in analogous experiments using AAV-Flex-TagRFP as the control virus (fig. S10). These results demonstrated that inhibiting the mPFC-vlPAG circuit disturbs placebo analgesia in SNI rats. Overall, suppressing MOR^+^ neuronal activity and activating the mPFC-vlPAG circuit may underlie placebo analgesia in SNI rats. Thus, we further investigated whether and how MOR^+^ neurons regulate the mPFC-vlPAG circuit.

### Monosynaptic connections from MOR^+^ neuron to vlPAG-projecting L5PN in the mPFC

First, we examined if there are direct monosynaptic connections between MOR^+^ neurons and L5PN in the mPFC that project into the vlPAG using a rabies virus-mediated retrograde trans-synaptic tracing method in MOR-Cre KI rats (*28*). Thus, we microinjected AAVrg-FLPo into the vlPAG region, followed by the microinjection of a mixture of AAV-fDIO-TCb-mCherry and AAV-fDIO-RG into the L5 mPFC in MOR-Cre KI rats. To identify MOR^+^ neurons, AAV-Flex-hM4Di-TagRFP was microinjected into the L5 mPFC. Two weeks later, EnvA-pseudotyped glycoprotein-deleted rabies viruses expressing GFP (EnvA+RVdG-GFP) were microinjected into the L5 mPFC to initiate trans-synaptic tracing (Fig. 3, A and B). The L5PN (Fig. 3Cb) that project into the vlPAG were identified by an antibody that can discriminate mCherry from TagRFP (Fig. 3Cc). Various GFP^+^ input neurons (Fig. 3Ca) were distributed in L5 and L2/3 of the mPFC, trans-synaptically labeled from starter neurons (Fig. 3Cd and Ch; yellow neurons) among L5PN (Fig. 3Cb). GFP^+^ input neurons co-expressed TagRFP but not mCherry (8±2.1%, four rats; Fig. 3C, f and h), indicating MOR^+^ neurons mono-synaptically connected with L5PN (Fig. 3C, e and h; blue and white neurons) that project into vlPAG. No mCherry or GFP expression was observed in rats without microinjection of AAVrg-FLPo into the vlPAG (fig. S11).

**Fig. 3.**
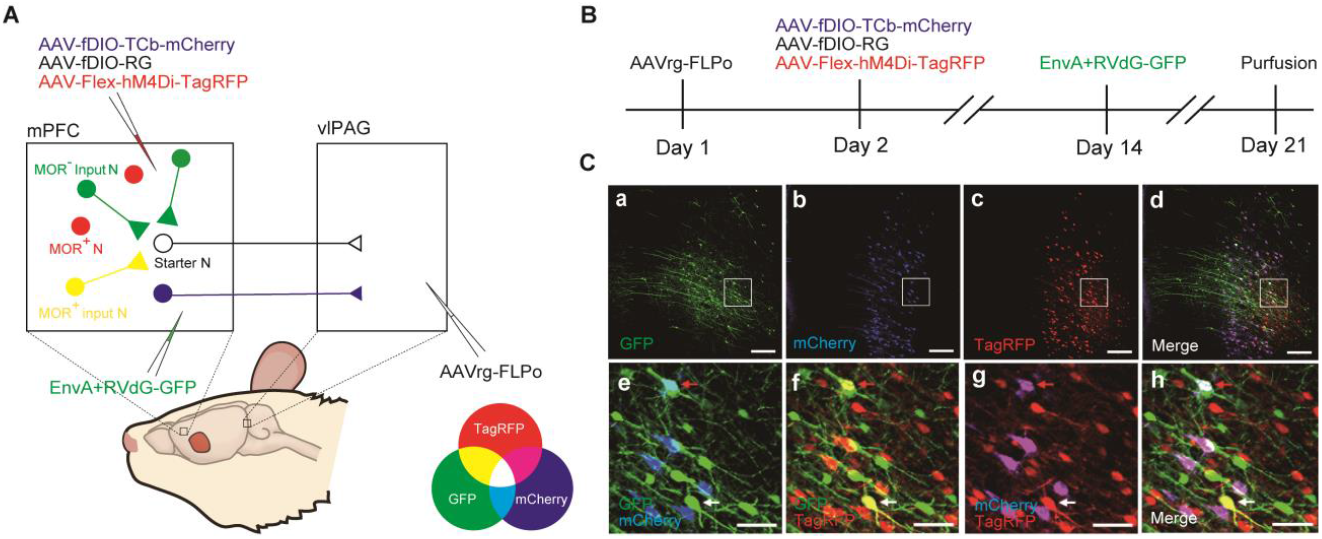
Monosynaptic connection between MOR^+^ neurons and mPFC-vlPAG circuit. (**A**) Schematic diagram for rabies virus-based retrograde trans-synaptic tracing. (**B**) Timeline of virus injection for rabies virus-based retrograde trans-synaptic tracing. (**C**) mPFC coronal sections show starter neurons labeled in white (41.3 ± 16.0, from four rats, expressed both GFP (green), mCherry (blue), and TagREP (red)); input neurons in green (165 ± 26.2, from four rats, GFP); projection neurons in magenta, merged by mCherry (blue) and TagRFP (red); and MOR^+^ neurons in red (TagRFP). Images of e-h in C indicate higher magnification of white rectangles in the images of a-d in (**C**). Note that neurons labeled in yellow (merged by GFP (green) and TagRFP (red)) in the images f and h indicate MOR^+^ neurons monosynaptically connected with projection neurons that project into the vlPAG. Scale bar: (**Ca to Cd**) 200 μm (**Ce to Ch**) 50 μm.

### MOR^+^ neurons predominantly inhibit excitatory outflow of mPFC to the vlPAG

Subsequently, we examined how MOR^+^ neurons regulate L5PN that project to the vlPAG using whole-cell patch-clamp recording in PFC slices from SNI rats 10 days after surgery. We concomitantly microinjected CTB647 into the vlPAG for retrograde labeling of L5PN in the mPFC and AAV-Flex-hChR2(H134R)-mCherry into the L5 of the mPFC to express excitatory opsin, ChR2(H134R), and mCherry in MOR^+^ neurons in MOR-Cre KI rats (Fig. 4, A and B).

**Fig. 4.**
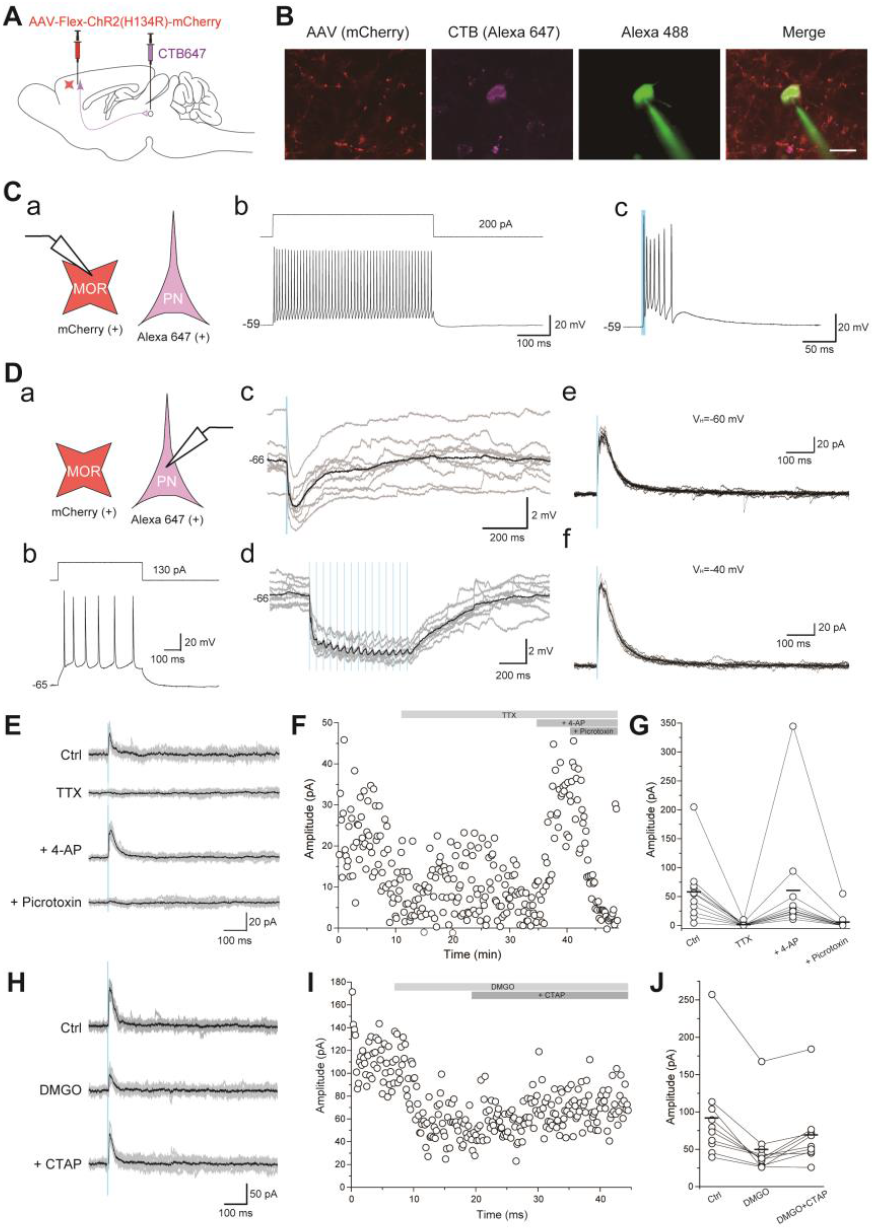
MOR^+^ neurons predominantly inhibit the excitatory outflow of mPFC to the vlPAG. (**A**) Schematic diagram for optogenetic manipulation of MOR^+^ neurons and labeling L5PN in the mPFC that project into the vlPAG. (**B**) Images show representative MOR^+^ neurons (red), L5PN (magenta, CTB647), and representative recording neurons (green, Alexa 488 contained in internal electrode solution). Scale bar: 20 μm. (**C**) Whole-cell patch-clamp recording from MOR^+^ neurons. a, Schematic diagram, b, Representative firing responses induced by current injection, and c, Representative firing responses induced by photostimulation. (**D**) Whole-cell patch-clamp recording from CTB647^+^L5PN that project into the vlPAG. a, Schematic diagram, b, Representative firing responses induced by current injection, c-d, Current-clamp recording shows photostimulation induced IPSP, e-f, Voltage-clamp recording shows photostimulation induced IPSC. (**E to G**) Voltage-clamp recording of IPSC form CTB647^+^L5PN without (Ctrl) and with TTX, or TTX and 4-AP, or TTX, 4-AP, and Picrotoxin. (**H to J**) Voltage-clamp recording of IPSC form CTB647^+^L5PN without (Ctrl) and with DAMGO or DAMGO and CTAP.

Most mCherry^+^ neurons (87.0%, 20/23) showed high-frequency action potentials without apparent frequency accommodation (Fig. 4C, a and c), suggesting they were fast-spiking interneurons. In the mCherry^+^ fast-spiking interneurons, single blue laser irradiation induced multiple action potentials (81.0%, 17/21; Fig. 4Cc) or excitatory postsynaptic potentials (19.0%, 4/21). CTB647^+^ neurons (28 neurons from 10 rats) showed a representative firing pattern of L5PN (Fig. 4D, a and b). Single blue laser irradiation induced inhibitory postsynaptic potentials (IPSPs) (0.5 ± 0.1 mV, n = 28). We further discovered that 15 train photostimulation induced IPSP summation followed by a sustained hyperpolarization: 1.8 ± 0.3 mV in amplitude and 528.7 ± 84.3 ms from the onset to peak (n = 28; Fig. 4, c and d). Under the voltage-clamp condition, inhibitory postsynaptic current (IPSC) amplitude was 8.3 ± 3.0 pA (n = 28) at the holding potential of -60 mV and 56.0 ± 12.4 pA at -40 mV (n = 27; Fig. 4, e and f). We then examined whether CTB647^+^ L5PN received monosynaptic inhibitory inputs from mCherry^+^ neurons (Fig. 4, E to G). Applying 1 μM tetrodotoxin abolished blue light-evoked IPSCs from 63.6 ± 17.8 pA to 1.7 ± 1.1 pA (n = 11, Fig. 4, E to G). 4-AP (1 mM) application recovered IPSCs to 64.7 ± 33.7 pA (n = 11; Fig. 4G), suggesting that photostimulation-evoked outward currents were mediated via monosynaptic connections. Outward currents were abolished by picrotoxin (100 μM; n = 11; Fig. 4G), indicating the involvement of GABA_A_ receptors. Bath application of DAMGO (1 μM), a MOR agonist, reduced the photostimulation-evoked IPSC amplitude from 92.0 ± 20.9 pA to 49.8 ± 14.1 pA, whereas CTAP (2 μM), a selective MOR antagonist, recovered the DAMGO-induced suppression of photostimulation-evoked IPSCs to 69.2 ± 14.4 pA (n = 10; Fig 4, H to J). These results suggested that MOR^+^ neurons directly inhibit L5PN, which project to the vlPAG via the GABA_A_ receptors.

### mPFC-vlPAG circuit is essential for MOR^+^ neuron-mediated pain modulations

Finally, to confirm whether the mPFC-vlPAG circuit is essential for MOR^+^ neuron-mediated pain modulation, we selectively ablated the L5PN in the mPFC that projected into the vlPAG and examined pain behavior in response to specific manipulation of MOR^+^ neuron activity. The selective ablation of mPFC-vlPAG was conducted using an immunotoxin (ITX; anti-Tac(Fv)-PE38, a gift from Prof. Ira Pastan)-mediated neural circuit elimination approach in which the AAVrg encoding human interleukin-2 receptor α-subunit (IL-2Rα) and EGFP (AAVrg-IL-2Rα-EGFP) was microinjected into the vlPAG to retrogradely express IL-2Rα in the mPFC L5PN, followed by the microinjection of recombinant ITX into the mPFC for selective ablation of the target neurons (Fig. 5, A to D, and F). Similar to our previous reports (*29*), the number of EGFP-expressing L5PN that project into the vlPAG was significantly reduced by the injection of ITX (5 ng/ul) compared with the injection of PBS (Fig. 5, E, G, and H). However, the number of TagRFP-expressing MOR^+^ neurons in the surrounding area was not changed in either ITX-or PBS-injected rats (Fig. 5, E, G, and I), indicating that the mPFC-vlPAG circuit was selectively ablated. The chemogenetic suppression of MOR^+^ neurons in these rats revealed decreased pain hypersensitivity in PBS-injected rats expressing AAV-Flex-hM4Di-TagRFP in MOR^+^ neurons in the mPFC (Fig. 5, J, K, and Fig. 1H). In contrast, this analgesic effect completely disappeared in ITX-injected rats (Fig. 5, J and K), indicating that the mPFC-vlPAG circuit is essential for MOR^+^ neuron-mediated pain modulation.

**Fig. 5.**
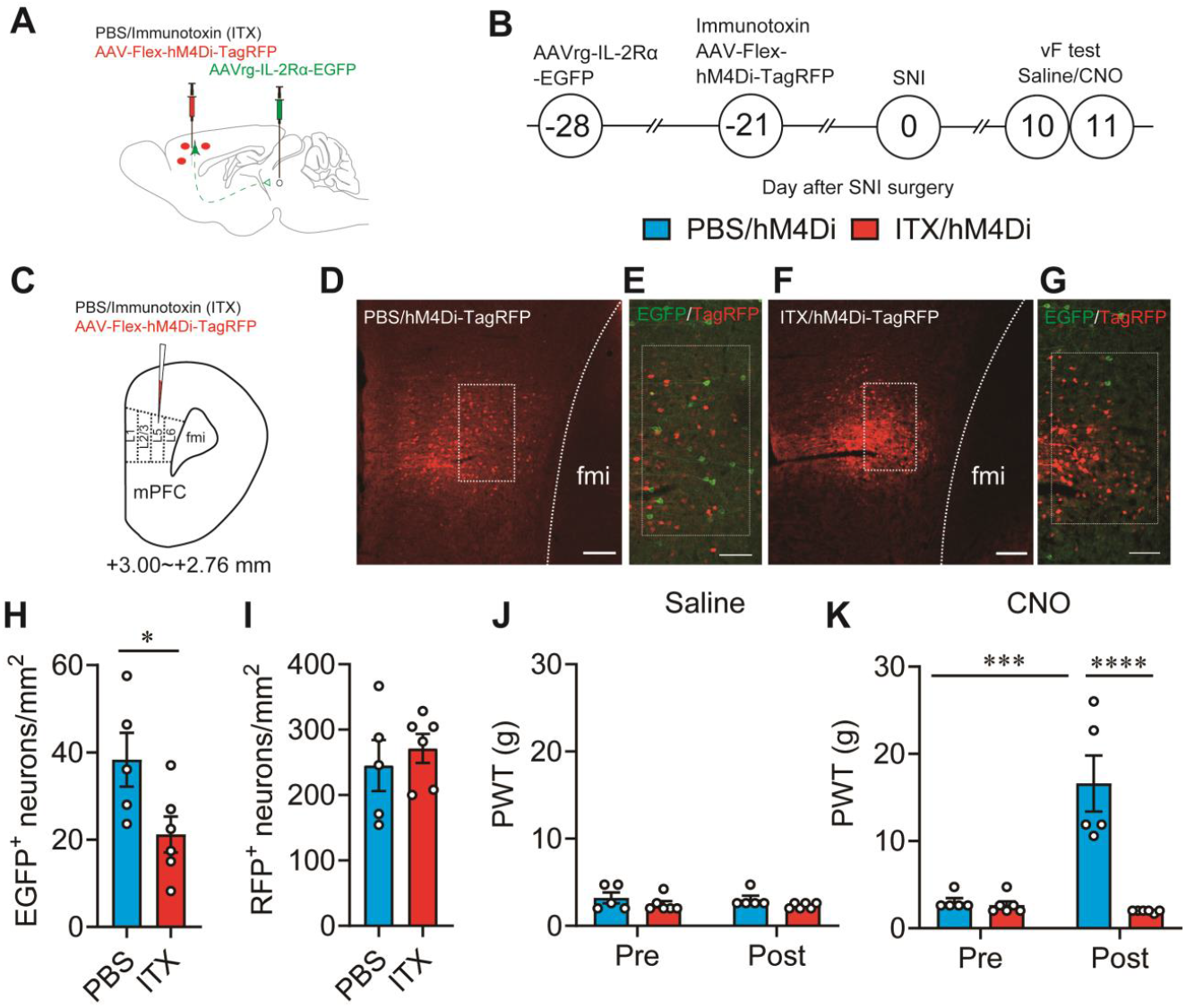
mPFC-vlPAG circuit is essential for MOR^+^ neuron-mediated pain modulations. (**A to C**) Schematic diagram for specific ablation of mPFC-vlPAG circuit using an immunotoxin-mediated elimination method and the timeline for chemogenetic manipulation and behavior test. (**D and F**) Images show TagPRF expression and counting area (white square) in the mPFC in PBS and ITX-treated rats. Scale bar: 200 μm. (**E and G**) High-magnified images of counting areas indicated by white squares in D and F showing TagRFP^+^ and GFP^+^ neurons. Scale bar: 100 μm. (**H**) The number of GFP^+^ neurons in PBS and ITX-treated groups. (**I**) The number of TagRFP^+^ neurons in PBS and ITX-treated groups. (**J and K**) Changes in PWT after saline (**J**) or CNO (**K**) injection in MOR-Cre KI rat with PBS/hM4Di (light blue column) or ITX/hM4Di (red column). \**P*<0.05, \*\*\**P*<0.001, \*\*\*\**P*<0.0001. All statistical information details were presented in supplementary table.

## Discussion

We demonstrated the fundamental neurobiological mechanisms underlying placebo analgesia. Intrinsic opioid signaling in the mPFC disinhibits excitatory outflow to the vlPAG via the suppression of MOR^+^ neurons, leading to the activation of the descending pain inhibitory system. We provide evidence that 1) chemogenetic activation of MOR^+^ neurons in the mPFC inhibited pharmacological conditioning-induced placebo analgesia in rats, 2) chemogenetic suppression of the mPFC-vlPAG circuit also blocked conditioning-induced placebo analgesia, and 3) MOR^+^ neurons in the mPFC were monosynaptically connected and directly inhibited L5PN that project to the vlPAG via the GABA_A_ receptors.

Placebo analgesia is evoked by an inert treatment that is inffective, suggesting that an endogenous pain inhibitory system must be activated by higher-order neuropsychological processes, such as expectations. Neuroimaging studies in humans have demonstrated that several prefrontal cortical regions, such as the rostral anterior cingulate cortex and dorsolateral PFC, are activated by placebo administration and subsequently recruit the lower parts of the descending pain inhibitory system involving the vlPAG and rostroventromedial medullar to suppress pain (*7, 30, 31*). The functional coupling between these prefrontal regions and the vlPAG is enhanced during placebo analgesia (*7*). The dorsolateral PFC in humans is one of the highest cortical regions involved in cognitive and affective components of pain modulation (*32*). Based on neuroanatomical characteristics and functional similarity, the mPFC in rodents is thought to be the homolog of the primate dorsolateral PFC (*33, 34*). In rodents, the mPFC regulates the descending pain inhibitory system via the vlPAG (*23, 25-27*). We have previously demonstrated that the functional coupling between the mPFC and vlPAG was increased in pharmacological conditioning-induced placebo analgesia in rats (*13*). These observations suggest that the current data can provide a fundamental mechanism of placebo analgesia and that the human placebo effect involves similar neurobiological processes.

Endogenous opioids interacting with MOR in hierarchical brain regions have been implicated in placebo analgesia, including the higher-order frontal cortex (*3, 8*). In humans, pharmacological functional magnetic resonance imaging and positron emission tomography imaging studies have demonstrated that opioid signals in multi-level brain regions such as dorsolateral PFC, rostral anterior cingulate cortex, PAG, and rostroventromedial medullar contribute to pain modulation and placebo analgesia (*7, 35, 36*). Behavioral pharmacological studies in animals have also demonstrated that placebo analgesia may be mediated by endogenous μ-opioid signals in the frontal cortical areas (*13, 37, 38*). However, the fundamental neuronal mechanisms of where and how the endogenous μ-opioid signal activates the descending pain modulation system have not been clarified. MOR, a major opioid receptor related to opioidergic analgesia, is broadly expressed from the peripheral nerves to higher-order brain regions and has been extensively discussed as a key player in pain modulation (*18, 36, 39-41*). In cerebral cortical regions, MOR is expressed in inhibitory GABAergic interneurons and modulates excitatory outflow to subcortical regions via the disinhibition of glutamatergic projection PN (*42*). We also confirmed that most MOR^+^ neurons around the L5 of the mPFC were GABAergic interneurons (Fig. 1D). Optogenetic suppression of GABAergic interneurons in the mPFC facilitates excitatory outflow to the vlPAG, producing an analgesic effect in an animal model of neuropathic pain (*25*). We demonstrated that MOR^+^ neurons monosynaptically connected with L5PN that project into the vlPAG and selective optogenetic activation of MOR^+^ neurons predominantly inhibit these neurons via GABA_A_ receptors (Fig. 4, E to G). Furthermore, we demonstrated that the chemogenetic activation or suppression of MOR^+^ neurons facilitates or inhibits pain behavior in neuropathic pain rats, respectively (Fig. 1, G and H). Moreover, we recently demonstrated that the microinfusion of a MOR antagonist into the mPFC suppresses placebo analgesia in rats (*13*). These observations suggest that endogenous μ-opioid signaling in the mPFC may suppress MOR^+^ GABAergic interneurons, leading to the disinhibition of excitatory outflow to the vlPAG mediated by the monosynaptic GABA_A_ receptor, which initiates placebo analgesia.

Placebo effects exist in any medical treatment and affect the therapeutic outcomes. The present study demonstrated the fundamental mechanisms of placebo analgesia that enable us to explore placebo effects in medical practice to maximize therapeutic efficacy and reduce adverse drug effects and tolerance. Opioidergic drugs are the first choice for alleviating cancer pain; however, several serious side effects, such as addiction, respiratory depression, and sedation, limit their clinical application. According to The Global Burden of Diseases, Injuries, and Risk Factors Study estimate, around 109,500 people died from opioid overdose in 2017 (*43*). Evidence-based clinical application of placebo effects may provide an effective solution for such socio-medical issues.

## Acknowledgments

We thank Editage for manuscript editing help.

## Funding

This work was supported by JSPS KAKENHI Grant Number JP 20K16511 to HN, 24659574, 26112003, and 15K14328 to YC, and by JP 16H06276 (AdAMS).

## Author contributions

Conceptualization: YC

Methodology: HN, YW, YN, SK, TS, TT, KM, TM, NM, YM, MK, KK, YC

Investigation: HN, YW, YN, TS, MS, MI, NM Visualization: HN, YW, YN, TS, NM Funding acquisition: HN, YC

Project administration: YW, MK, KK, YC Supervision: YC

Writing – original draft: HN, YW, YN, TS, NM, MK, YC

Writing – review & editing: HN, YW, YN, TS, TT, KM, MK, YW, KK, YC

## Competing interests

The authors declare no competing interests.

### Data and materials availability

All data are available in the main text or the supplementary materials.

## Supplementary Materials

### Materials and Methods

Figs. S1 to S11

Tables S1 to S4

References (*1*–*13*)

## Notes

### Competing Interest Statement

The authors have declared no competing interest.

