## Supplemental Files for "Opioidergic activation of descending pain inhibitory system underlies placebo analgesia"

**This PDF file includes:**

Materials and Methods
Figs. S1 to S11
Tables S1 to S4
References (*1-13*)

### Materials and Methods

#### Animals

All experiments in this study were approved by the Animal Research Committee of Osaka University (Permission number: 24-006-042), the Institutional Animal Care and Use Committee (IACUC) of RIKEN, Kobe, and Nihon University. Iar: Long–Evans rats for donor of embryos were obtained from Japan SLC Inc. (Shizuoka, Japan). Iar: Wistar–Imamichi rats for transplant recipients of genome-edited zygotes were obtained from the Institute for Animal Reproduction, Ibaraki, Japan. All animals were housed at 23±1.5°C temperature, 45±15% humidity, and a 12:12 h light:dark cycle. They were fed a standard pellet diet (Oriental Yeast Co., Tokyo, Japan) and tap water *ad libitum*. Behavioral experiments were performed following the Principles of Laboratory Animal Care (NIH Publication No. 85-23, revised 1985). Behavioral studies were performed according to the Animal Research Reporting of In Vivo Experiments (ARRIVE) guidelines (*1*). All efforts were made to minimize animals used and their suffering.

#### Generation of μ-opioid receptor (MOR)-Cre knock-in (KI) rats by CRISPR/Cas9 genome editing

To selective functional manipulation of μ opioid receptor-positive ($MOR^+$) neuron activity in vivo, we designed and developed genetically engineered rats expressing Cre recombinase under the control of the MOR 1 (*Oprm1)* promoter using CRISPR/Cas9 genome editing technology (fig. S1A).

*1) Preparation of CRISPR components and long single-stranded donor DNA*

The gRNA was designed using CRISPOR software (http://crispor.tefor.net/), which predicts unique target sites throughout the rat genome. The target sequence selected for KI production is 5’-GTGAGACCCAGTTAGGGCAA-3’. gRNA was prepared using a Precision gRNA Synthesis Kit (Thermo Fisher, Waltham, MA, USA). Cas9 mRNA was transcribed *in vitro* using the mMESSAGE mMACHINE T7 Ultra Kit (Thermo Fisher) from a linearized plasmid (ID #72602; www.addgene.org/CRISPR) and purified using a MEGAClear kit (Thermo Fisher). As a KI donor, long single-stranded DNA (lssDNA) was prepared using nicking endonucleases (nickases), as reported previously (*2*). The lssDNA introduced into fertilized embryos consisted of the left homology arm, T2A, nlsCre, polyA signal, and the right homology arm sequences in order (fig.S1A).

*2) Manipulation of rat embryos and electroporation*

For electroporation, 100 Iar: Long-Evans embryos at 6–8 h after collection were placed into a chamber with 40 μL of serum-free medium (Gibco Opti-MEM, Thermo Fisher Scientific) containing 400 ng/μL Cas9 mRNA, 200 ng/μL gRNA, and 20 ng/μL lssDNA. They were electroporated with a 5 mm gap electrode (CUY520P5 Nepa Gene, Chiba, Japan) in a NEPA21 Super Electroporator (Nepa Gene, Chiba, Japan) (*3*).

Seventy-two embryos that developed to the two-cell stage after Cas9 mRNA, gRNA, and lssDNA introduction were transferred into the oviducts of 3 female surrogates anesthetized with isoflurane (DS Pharma Animal Health Co., Ltd., Osaka, Japan). Finally, three pups were born, one of which was confirmed to be a 2A-nlsCre KI rat (Table S1).

*3) Genotyping analysis and sequencing for MOR-Cre KI rat*

Genotyping polymerase chain reaction (PCR) was performed using genomic DNA extracted from the tail as a template and KAPA2G Fast Hotstart ReadyMix with dye (KK5610, KAPA BIOSYSTEMS, MA, USA). The PCR products amplified with specific primer sets (Table S2) were directly sequenced using the BigDye terminator v3.1 cycle sequencing mix and the standard protocol for an Applied Biosystems 3130 DNA Sequencer (Life Technologies, CA, USA). The PCR analysis validated the correct fragment size and sequence for the internal 5' and 3' junction of the transgene insertion into the target locus (fig. S1B). Homozygous MOR-Cre KI rats were born at the expected Mendelian ratios and were viable and fertile without noticeable gross abnormalities. MOR-Cre KI rats are available to the scientific community upon request.

**Virus vector production**

All the viral vectors are presented in Table S3. Adeno-associated virus (AAV)5-CaMKIIα-EGFP (AAV-CaMKIIα-EGFP), AAV2-hSyn-DIO-hM3Dq-mCherry (AAV-DIO-hM3Dq-mCherry), retrograde AAV (AAVrg)-pgk-Cre (AAVrg-Cre), AAVrg-hSyn-EGFP (AAVrg-EGFP) and were purchased from Addgene (MA, USA). AAV5-CAG-FLEx (FRT)-TC-mCherry (AAV-fDIO-TCb-mCherry), and AAV8-CAG-FLEx (FRT)-G (AAV-fDIO-RG) were generated in the UNC vector core using the plasmids (Addgene #67827 and #67828). AAV2-EF1α-Flex-hM3Dq-2A-cgfTagRFP (AAV-Flex-hM3Dq-TagRFP), AAV2-EF1α-Flex-hM4Di-2A-cgfTagRFP (AAV-Flex-hM4Di-TagRFP), AAVrg-CAGGS-IL-2Rα-EGFP (AAVrg- IL-2Rα-EGFP), AAV2-EF1α-Flex-ChR2(H134R)-mCherry (36) (AAV-Flex-ChR2(H134R)-mCherry), AAVrg-EF1a-FLPo (AAVrg-FLPo) were obtained from our laboratory. These AAVs were prepared using an AAV Helper-Free System (Agilent Technologies, CA, USA) (*4*). HEK293T cells were transfected with the transfer gene, replication/encapsulation gene, and adeno-helper gene plasmids using the calcium phosphate precipitation method. The two rounds of CsCl gradient ultracentrifugation at 100,000 × *g* and 4°C for 23 h. The fractions containing AAV were collected and subjected to three rounds of dialysis, and the dialyzed solution was concentrated by centrifugation through a Vivaspin Turbo filter (Sartorius, Gottingen, Germany). The viral genome titer of the AAV vectors was determined by quantitative PCR using the TaqMan System (Thermo Fisher Scientific). To prepare Rabies dG-GFP+EnvA, we unitized the viruses and cell lines, as described previously (*5*). RVdG-GFP was de novo prepared using the B7GG cell line (a gift from Ed Callaway) and plasmids (pCAG-B19N, pCAG-B19P, pCAG-B19L, pCAG-B19G, and pSADdG-GFP-F2; gifts from Ed Callaway), and then pseudotyped using BHK-EnvA cells (a gift from Ed Callaway). The titer of RVdG-GFP+EnvA used in this study was estimated to be $4 \times 10^9$ infectious particles per mL based on serial dilutions of the virus stock, followed by infection of the HEK293-TVA800 cell line (a gift from Ed Callaway).

**Stereotaxic microinjection**

Rats were anesthetized with 4–5% isoflurane, and anesthesia was kept with 1.5% isoflurane using a Small Animal Anesthetizer (NS-AN-20, NEUROSCIENCE, Inc., Tokyo, Japan). The head of each rat was fixed in a small animal stereotaxic instrument with a Digital Display Console (Model 68025, David Kopf, CA, USA). The skull was exposed through an incision in the head of a rat using a surgical knife. A small burr hole was drilled for microinjection using an Ideal Micro-Drill ™ (CellPoint Scientific Inc., MD, USA). After carefully removing the dura mater, neural tracers, and AAVs (Table S3), they were injected into each position using a UMP3 pump regulated by Micro-4 (flow rate: 20–50 nL/min; World Precision Instruments, FL, USA). After microinjection, the incision was sutured, an antimicrobial was injected to prevent infection, and the animal was returned to the home cage.

*1) Retrograde and anterograde labeling*

For retrograde somatic labeling (Fig. 2), Cholera Toxin Subunit B (recombinant) and Alexa Fluo$^{TM}$ 647 conjugate (CTB647; Thermo Fisher Scientific) were dissolved in phosphate-buffered saline (PBS) at a concentration of 1 mg/ml and microinjected into the ventrolateral periaqueductal gray (vlPAG; 200 nL). For anterograde tracing, 400 nL of AAV-CaMKIIα-EGFP was microinjected into layer five (L5) of medial prefrontal cortex (mPFC). The following coordinates were used: mPFC: AP +3.72, ML +0.7, DV +3.2; and vlPAG: AP -7.52, ML +0.65, DV +5.0.

*2) Monosynaptic retrograde rabies virus tracing*

To investigate whether there are direct monosynaptic connections between MOR$^{+}$ neurons in the mPFC and L5 pyramidal neurons (L5PNs) that project into the vlPAG (Fig. 3), 200 nL retrograde AAVrg-FLPo was injected into the vlPAG, and 600 nL of a 1:1 mixture of AAV-fDIO-TCb-mCherry and AAV-fDIO-RG was injected into the mPFC. To label MOR$^{+}$ neurons, 400 nL of AAV-Flex-hM4Di-TagRFP was injected into the mPFC. Two weeks after AAVs injection, 200 nL of Rabies dG-GFP+EnvA was injected into the mPFC. One week later, rats were perfused with 4% paraformaldehyde, and their brains were excised for further histological analysis. In the negative control (fig. S11), AAVs microinjection was performed analogously, except that AAVrg-FLPo was injected into the vlPAG.

*3) Selective ablation of mPFC-vlPAG pathway*

We also performed an immunotoxin-mediated neural circuit elimination approach for selective ablation of the mPFC-vlPAG pathway (Fig. 5). For this purpose, 1200 nL of a mixture of AAV-Flex-hM4Di-TagRFP (900 nL) and immunotoxin (100 nL, final 50 ng/μL). AAVrg-IL-2Rα-

EGFP (200 nL) was injected into vlPAG. The following coordinates were used: mPFC, AP +3.24/+3.00/+2.76, ML +0.7, DV +3.0; vlPAG, AP -7.56, ML +0.6, DV +4.8.

*4) optogenetic/chemogenetic manipulation*

For chemogenetic manipulation of $MOR^+$ neurons in the mPFC (Fig. 1 and fig. S7), 600 nL AAV-Flex-hM3Dq-2A-TagRFP, or 1200 nL AAV-Flex-hM4Di-TagRFP were injected into L5 of the mPFC. For chemogenetic manipulation of the mPFC-vlPAG circuit (Fig. 2, fig. S9 and 10), 300 nL of a mixture of AAVrg-Cre (500 nL) and CTB647 (200 nL) was injected into the vlPAG, and 1200 nL AAV-DIO-hM3Dq-mCherry or 1200 nL AAV-Flex-hM4Di-TagRFP was injected into L5 of the mPFC.

For optogenetic activation of $MOR^+$ neurons (Fig. 4), 1000 nL of AAV-Flex-ChR2(H134R)-mCherry was microinjected into the L5 of the mPFC. The following coordinates were used: mPFC, AP +3.00, ML ±0.7, and DV +2.7/+3.4.

**Spared nerve injury (SNI) model**

Neuropathic pain was induced by SNI of the unilateral sciatic nerve, as described previously (*6*). Briefly, rats (8–12 weeks) were anesthetized with 1.5% isoflurane, and a skin and muscle incision was made on the left hind limb to expose the sciatic nerve and its three branches. For SNI treatment, the common peroneal and tibial nerves were ligated, and distal to the trifurcation was cut, leaving the sural nerve intact. After surgery, the incision was sutured, an antimicrobial was injected to prevent infection, and the animal was then returned to the home cage. In SNI rats, the receptive field of the lateral aspect of the hind paw skin innervated by the sural nerve displays hypersensitivity to tactile stimuli (fig. S5).

**Measurements of pain threshold**

The pain threshold in rats with neuropathic pain was measured by the up-down method using von Frey filaments, as described previously (*7*). Briefly, rats were placed in a plastic cage with a wire mesh bottom and acclimatized for 30-60 min before testing. Stimulation of the bilateral hind paws was gradually induced by von Frey filaments (Aesthesio®, DanMic Global, LLC, CA, USA) from 2 to 26 g (2, 4, 6, 8, 15, and 26 g). Stimulation was presented at intervals of at least 30 s, and the pain response was determined when rats represented lifting, shaking, or biting the hind paw. Measurements of the pain threshold were started at least five days after SNI surgery. The paw withdrawal threshold is reportedly calculated based on pain score, which is acquired by pain response pattern, as 50% paw withdrawal threshold (*7*).

**Establishment of placebo analgesia in SNI rats and chemogenetic manipulation**

Placebo analgesia in rats with neuropathic pain was induced by pharmacological conditioning, as described previously (*8*). A commercially available painkiller, gabapentin hydrochloride (GBP; 100 mg/kg, Tokyo Chemical International, Tokyo, Japan), was used for pharmacological conditioning. As shown in fig. S5C, the analgesic effect of GBP was significantly observed 1 h after intraperitoneal injection and then decreased over time. The analgesic effect of single intraperitoneal injection of GBP was completely disappeared 24 h after the injection (fig. S5C). The pharmacological conditioning was established by pairing of GBP with the intraperitoneal injection (conditioned stimulus) in SNI rats for four consecutive days. During the conditioning period, the analgesic effects of GBP were assessed using the von Frey filament test. Placebo analgesia was evaluated by measuring the pain threshold after an intraperitoneal saline injection as a placebo on the test day (Day 5). In the chemogenetic manipulation, Clozapine-*N*-oxide

(CNO; 1 mg/kg, Enzo Life Science Inc., BML-NS105, NY, USA) was injected instead of saline on the test day for chemogenetic activation or suppression of specific neuronal activity.

**Histological experiments**

Under deep anesthesia with isoflurane, rats were transcardially perfused with PBS, followed by 4% paraformaldehyde (PFA, Nacalai, Osaka, Japan) in PBS. Then the brains were excised and soaked in 4% PFA to postfix overnight at 4°C. Serial brain sections were cut at 30 μm thickness using a cryostat (Cryostar NX70, Epredia Holdings Ltd., NH, USA) and collected in dishes containing PBS with 0.1% Sodium Azide (FUJIFILM Wako Pure Chemical Corporation, Osaka, Japan). For in situ hybridization (ISH), the brain sections were mounted onto MAS coat slide glass (MATSUNAMI, Osaka, Japan) and kept in -30°C freezer.

*1)Immunohistochemistry*

For immunostaining, brain sections were incubated overnight at 4°C with the following primary antibodies respectively: rabbit anti-RFP (R10367; 1:1000; Thermo Fisher Scientific), mouse anti-c-Fos (ab208942; 1:500, Abcam, Cambridge, UK), rabbit anti-mCherry (ab167453; 1:1000, Abcam), chicken anti-GFP (ab13970, 1:1000, Abcam), rabbit anti-GFP (A6455; 1:1000, Thermo Fisher Scientific). The next day, brain sections were washed with 0.3% Triton X-100 in PBS and were incubated with secondary antibodies, Alexa Fluor™ conjugated secondary antibodies (1:500, Thermo Fisher Scientific), or Jackson antibodies (1:1000, Jackson ImmunoResearch Laboratories Inc., PA, USA) for 1 h at RT. Brain sections were mounted on APS glass slides (MATSUNAMI, Osaka, Japan) with ProLong™ Gold Antifade Mountant (Thermo Fisher

Scientific) and 4',6-Diamidino-2-phenylindole (DAPI) (5 ng/mL; 340-07971, DOJINDO, Kumamoto, Japan).

*2) RNAscope® in situ hybridization*

In situ hybridization with RNAscope® was performed according to the RNAscope® Multiple Fluorescent Reagent Kit v2 Assay User manual for Fresh Frozen sections (Advanced Cell Diagnostics, Bio-Techne Corporation [ACD], CA, USA, #323100).

Probes targeting intronic regions for solute carrier family 32 members 1*, Slc32a1* (VGAT; ACD Cat #424541, accession number: NM_031782.1, probe region 288-1666), *Slc17a6* (vGlut2; ACD Cat #317011, accession number: NM_053427.1, probe region 1109-2024), *Cre* (ACD Cat #312281), and *Tag-RFP* (ACD # 1190361) were custom-designed and synthesized. *Cre* and *Tag-RFP* were labeled with TSA Cyanine 3 (Perkin Elmer # NEL744001KT, 1:1000 dilution), *and Slc32a1* was labeled with TSA Fluorescein (Perkin Elmer # NEL741001KT, 1:1000 dilution). Fixed sections were pretreated with hydrogen peroxide for 10 min and Protease III for 30 min, and the probes were hybridized and amplified according to the manufacturer's instructions.

*3) Acquisition and analyzed fluorescence images data*

Fluorescent images were acquired using a Zeiss LSM 710 confocal microscope (Carl Zeiss, Oberkochen, Germany) and analyzed using ZEN software (Carl Zeiss). In Fig 5, the counting area of layer 5 is defined as in previous reports (*9, 10*).

**Electrophysiology**

*1) Whole-cell patch-clamp*

For optogenetic activation of $MOR^+$ neurons in the mPFC, 500 nL of AAV-ChR2(H134R)-mCherry was microinjected into the layer 5 of the mPFC of MOR-Cre KI rats. To identify pyramidal neurons in the mPFC that projected into the vlPAG, we injected a retrograde tracer, CTB647 (200 nL), into the vlPAG. Subsequently, 1–2 weeks after AAV injection, the animals were used to prepare histology specimens as described previously (*11, 12*) except for the recipe of ice-cold modified artificial cerebrospinal fluid (M-ACSF) (*13*). Briefly, rats were deeply anesthetized with isoflurane (5%) and perfused transcardially with the following ice-cold M-ACSF: 2.5 mM KCl, 0.5 mM $CaCl_2$, 10 mM $MgSO_4$, 1.25 mM $NaH_2PO_4$, 2 mM thiourea, 3 mM sodium pyruvate, 92 mM N-methyl-D-glucamine, 20 mM N-(2-hydroxyethyl)piperazine-N'-2-ethanesulfonic acid (HEPES), 25 mM D-glucose, 5 mM L-ascorbic acid, and 30 mM $NaHCO_3$ (pH 7.35–7.40). After decapitation, the tissue blocks, including the mPFC, were rapidly removed and stored in ice-cold M-ACSF. Coronal slices were cut at a thickness of 350 μm using a microslicer (Linearslicer Pro 7, Dosaka EM, Kyoto, Japan) and were incubated in the holding ACSF (H-ACSF) at 32 °C; the composition of H-ACSF was 2.5 mM KCl, 2 mM $CaCl_2$, 2 mM $MgSO_4$, 1.25 mM $NaH_2PO_4$, 2 mM thiourea, 3 mM sodium pyruvate, 92 mM NaCl, 20 mM HEPES, 25 mM D-glucose, 5 mM L-ascorbic acid, and 30 mM $NaHCO_3$ (pH 7.35–7.40). The slices were then transferred to a recording chamber containing normal ACSF (N-ACSF) at room temperature. The N-ACSF recipes were 126 mM NaCl, 3 mM KCl, 2 $MgSO_4$, 1.25 mM $NaH_2PO_4$, 26 mM $NaHCO_3$, 2 mM $CaCl_2$, and 10 D-glucose (in mM) at room temperature. All ACSF, M-, H-, and N-ACSF were continuously aerated with a mixture of 95% $O_2$ and 5% $CO_2$. To visualize $MOR^+$ and $CTB647^+$ neurons, these slices were transferred to a recording chamber on a confocal microscope (FV-1000, Olympus, Tokyo, Japan), and the injection sites were imaged through a 60 × objective with a resolution of 1600 × 1600 pixels.

For recording, the slices, including the mPFC, were placed in a recording chamber perfused continuously with N-ACSF at a rate of 2.4 mL/min. To minimize drug adherence to the perfusion route, we used an infusion pump with a head made of polytetrafluoroethylene (Q-100-TT-P-S, TACMINA, Osaka, Japan) and silicon tubes (C-flex tubing, Cole-Parmer Instrument, IL, USA).

Thin-wall borosilicate patch electrodes (3–5 MΩ) were pulled using a Flaming-Brown micropipette puller (P-97, Sutter Instruments, CA, USA). The pipette solution contained the following components ($E_{Cl^-}$ = -65 mV): 135 mM potassium gluconate, 10 mM HEPES, 0.5 mM EGTA, 2 mM $MgCl_2$, 2 mM magnesium adenosine triphosphate (ATP), and 0.3 mM sodium guanosine triphosphate (GTP). The pipette solution had a pH of 7.3 and an osmolarity of 300 mOsm. The liquid junction potential of the pipette solution described above was -9 mV. Voltage was not corrected in this study.

The recordings were obtained at room temperature. The seal resistance was > 10 GΩ, and only data obtained from electrodes with access resistance of 6–20 MΩ and with < 20% change during recordings were included in this study. Alexa Fluor™ 488 (Thermo Fisher Scientific) was added to the internal solution in a subset of the experiments to identify the neural subtypes of the recorded neurons (Fig. 4).

Whole-cell patch-clamp recordings were obtained from $mCherry^+$ or $CTB^+$ neurons located in layer 5 using a fluorescence microscope equipped with Nomarski optics (BX61W1; Olympus, Tokyo, Japan), and an infrared-sensitive video camera (IR-1000; DAGE-MTI, IN, USA). Electrical signals were recorded with an amplifier (Multiclamp 700B, Molecular Devices, CA, USA) and a digitizer (Axon Digidata® 1440A, Molecular Devices), observed online, and stored on a computer hard disk using Clampex (pClamp 10, Molecular Devices).

The voltage responses of mCherry$^+$ or CTB$^+$ neurons were recorded by injecting depolarizing and hyperpolarizing current pulses (500 ms) to examine basic membrane properties, including input resistance, single spike kinetics, voltage-current relationship, and repetitive firing patterns and frequency.

To examine the photostimulation-evoked synaptic responses, ChR2, expressed in MOR$^+$ neurons, was activated by collimated blue light (470 nm) using an LED system (8.6 mW) through a water-immersion 40 × microscope objective. Photostimulation was applied to the slices with a 1–5 ms duration. We first recorded synaptic responses to single and 15-train photostimulation under the current clamp condition. Next, photostimulation-evoked synaptic responses were recorded under voltage-clamp conditions, in which the holding potentials were set at -60 mV or -40 mV.

To determine whether the photostimulation-evoked synaptic responses were monosynaptic or polysynaptic, we applied tetrodotoxin (TTX; 1 μM; Abcam), 4-aminopyridine (4-AP; 1 mM; Nacalai Tesque, Kyoto, Japan), and picrotoxin (100 μM; Sigma-Aldrich). To examine the effects of activating μ-opioid receptors, we administered DAMGO, a μ-opioid receptor agonist, with or without CTAP, a selective μ-opioid receptor antagonist. All drugs were bath applied to the perfusate. The other compounds were purchased from Wako Pure Chemical Industries. All recorded data were analyzed using Clampfit (pClamp 10, Molecular Devices). The average amplitude was obtained from 10–20 consecutive sweeps.

*2) In vivo extracellular multi-channel recordings*

To confirm the chemogenetic activation of MOR$^+$ neurons, we also recorded the extracellular action potential in the mPFC of MOR-Cre KI rats in which AAV-Flex-hM3Dq-TagRFP was microinjected into L5 of the mPFC (fig. S6). At least 2 weeks after the AAV injection, the rats were anesthetized with 1.8% isoflurane, and the heads of a rat were fixed in a stereotaxic instrument (Model 68025, David Kopf, CA, USA). A 32-channel silicon probe (A4 × 8–5mm-100-200-177, with eight electrodes in a linear arrangement on four shanks, NeuroNexus Technologies, Ann Arbor, MI, USA) was inserted vertically into the mPFC using a micromanipulator on a stereotaxic frame. The silicon probe was left in place for 30 min to allow the brain tissue to recover from the acute damage. Then, a 10-minute extracellular multi-channel recording of action potential was performed before and after CNO injection (1.0 mg/kg, i.p.). The 10 minutes of recordings were repeated every 30 min until 160 min after the CNO injection. Extracellular signals were amplified and filtered using a 32-channel head stage (band pass:0.3-7.5 kHz, Cereplex M, Blackrock Microsystems, Salt Lake City, USA). Amplified signals were digitized at 30 kHz using a head stage. Extracellular signals were processed offline to isolate spike events of individual neurons using the spike-sorting software BOSS (Blackrock Microsystems, US).

**Statistical analysis**

All data were calculated by the Graphpad Prism version 8 (Graphpad Software, San Diego, CA, USA) and shown as mean ± standard of mean (S.E.M). We used the two-way repeated measures (RM) ANOVA followed by Bonferroni's comparisons test, one-way ANOVA followed by Tukey's multiple comparisons test, unpaired t-test, and unpaired t-test with Welch's correction. Differences with a p-value of less than 0.05 were judged as statistically significant. All Statistical information were presented in supplemental table.

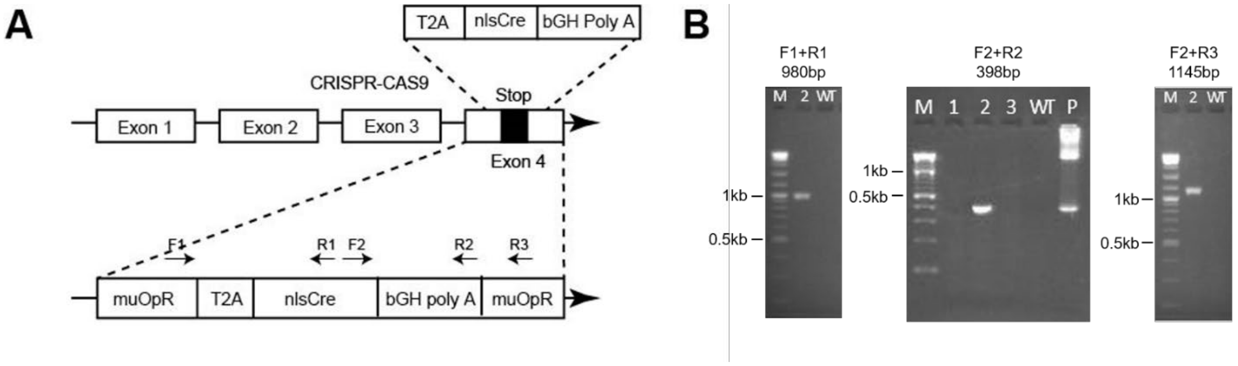


**Fig. S1. Construction of MOR-Cre KI rat.** (**A**) Targeting strategy for inserting the nlsCre. (**B**) Gel images for genotyping with electrophoresis of gene extracted from MOR-Cre KI rat.

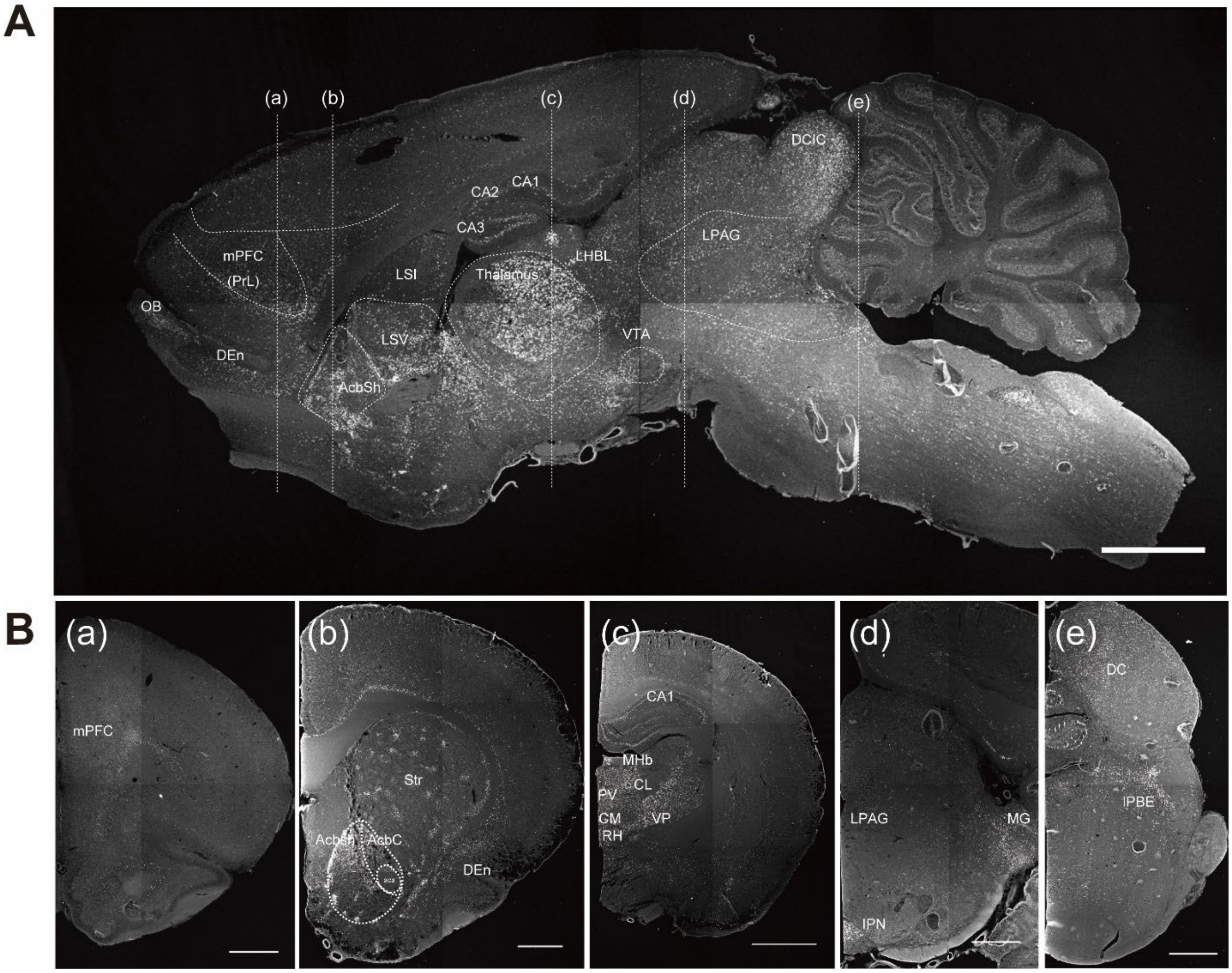


**Fig. S2. Distribution of MOR gene expression by RNAscope®.** (**A**) Distribution of MOR gene expression in sagittal brain section. (**Ba**) MOR gene expression in coronal brain section in cut surface (a) of A, including a mPFC of prelimbic cortex (PrL) (**Bb**) MOR gene expression in coronal brain section in cut surface (b) of A, including stratum (Str), nucleus accumbens (NAc), and DEn (**Bc**) MOR gene expression in coronal brain section in cut surface (c) of A, including the thalamus, habenula (Hb), and hippocampus (**Bd**) MOR gene expression in coronal brain section in cut surface (d) of A, including periaqueductal gray (PAG) and Intrapeduncular nucleus (IPN) (**Be**) MOR gene expression in coronal brain section in cut surface (e) of A, concluding DC and parabrachial nucleus (PB). Scale bar: (**A**) 2000 µm (**Ba**) 1000 µm (**Bb**) 1000 µm (**Bc**) 2000 µm (**Bd**) 1000 µm (**Be**) 1000 µm.

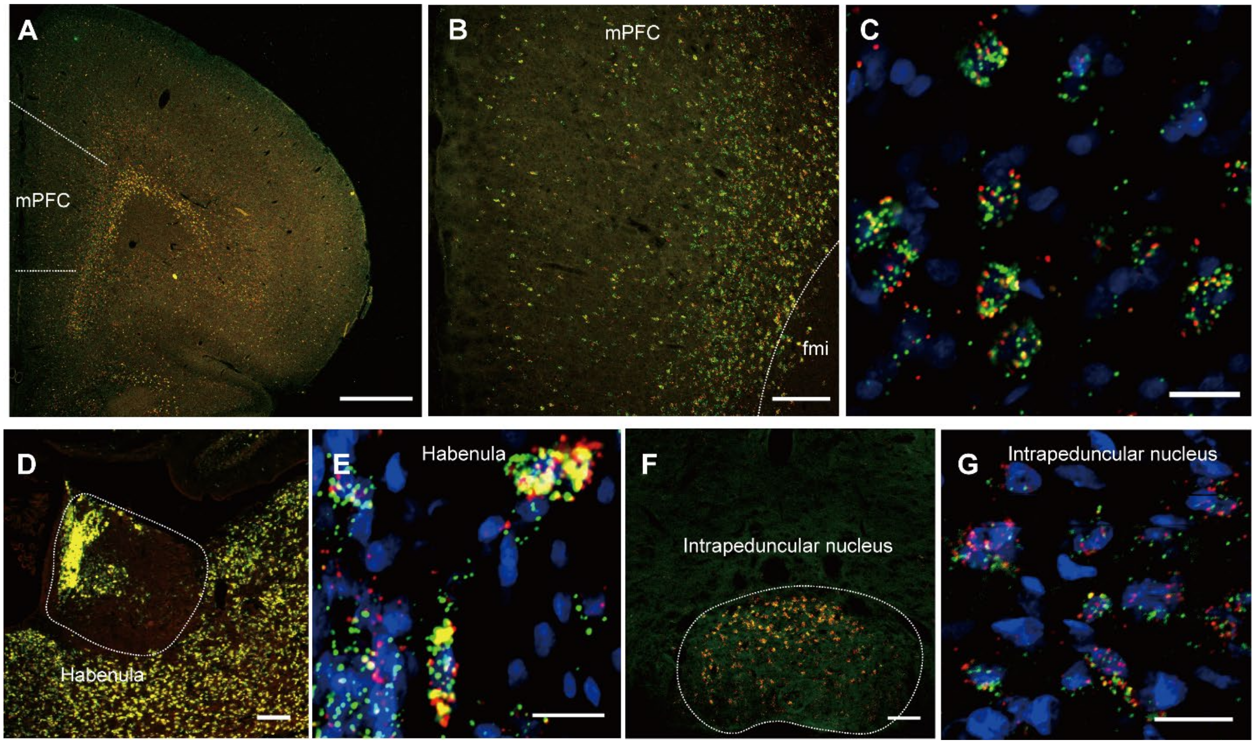


**Fig. S3. MOR specific expression of Cre gene in MOR-Cre KI rat.** (**A**) RNAscope® fluorescent images of co-express with MOR and Cre in mPFC (**B**) RNAscope® fluorescent images of Co-express with MOR and Cre in Habenula and IPN. Scale bar: (**A**) 1000 μm (**B**) 200 μm (**C**) 20 μm (**D**) 200 μm (**E**) 20 μm (**F**) 100 μm (**G**) 20 μm.

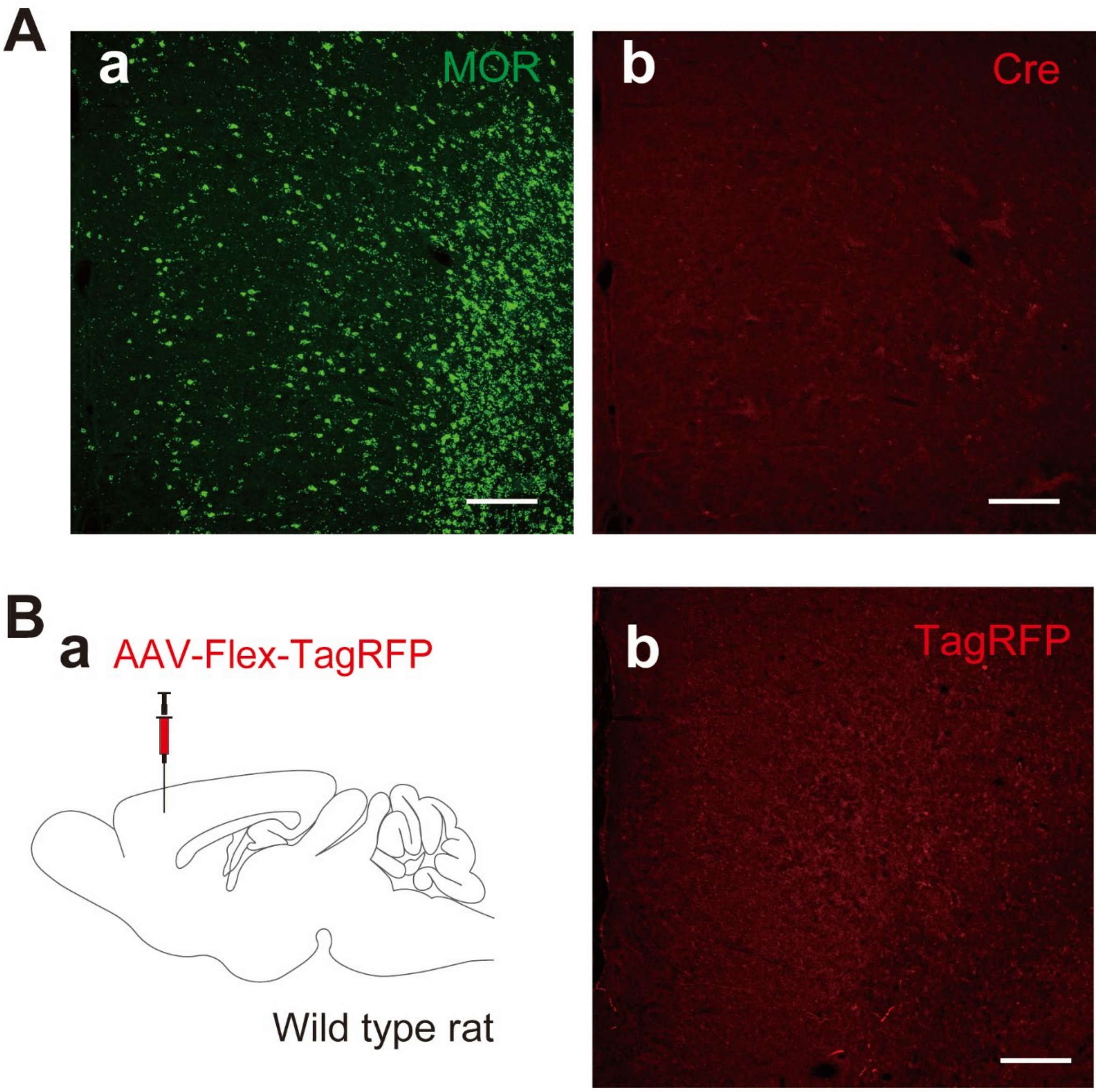


**Fig. S4. MOR specific expression of Cre gene in MOR-Cre KI rat.** (**A**) No expression of Cre gene by RNAscope® in wild type rats. (**B**) Immunostaining image of TagRFP in wild type rats injected AAV-Flex-hM4Di-TagRFP into mPFC. Scale bar: 200 μm

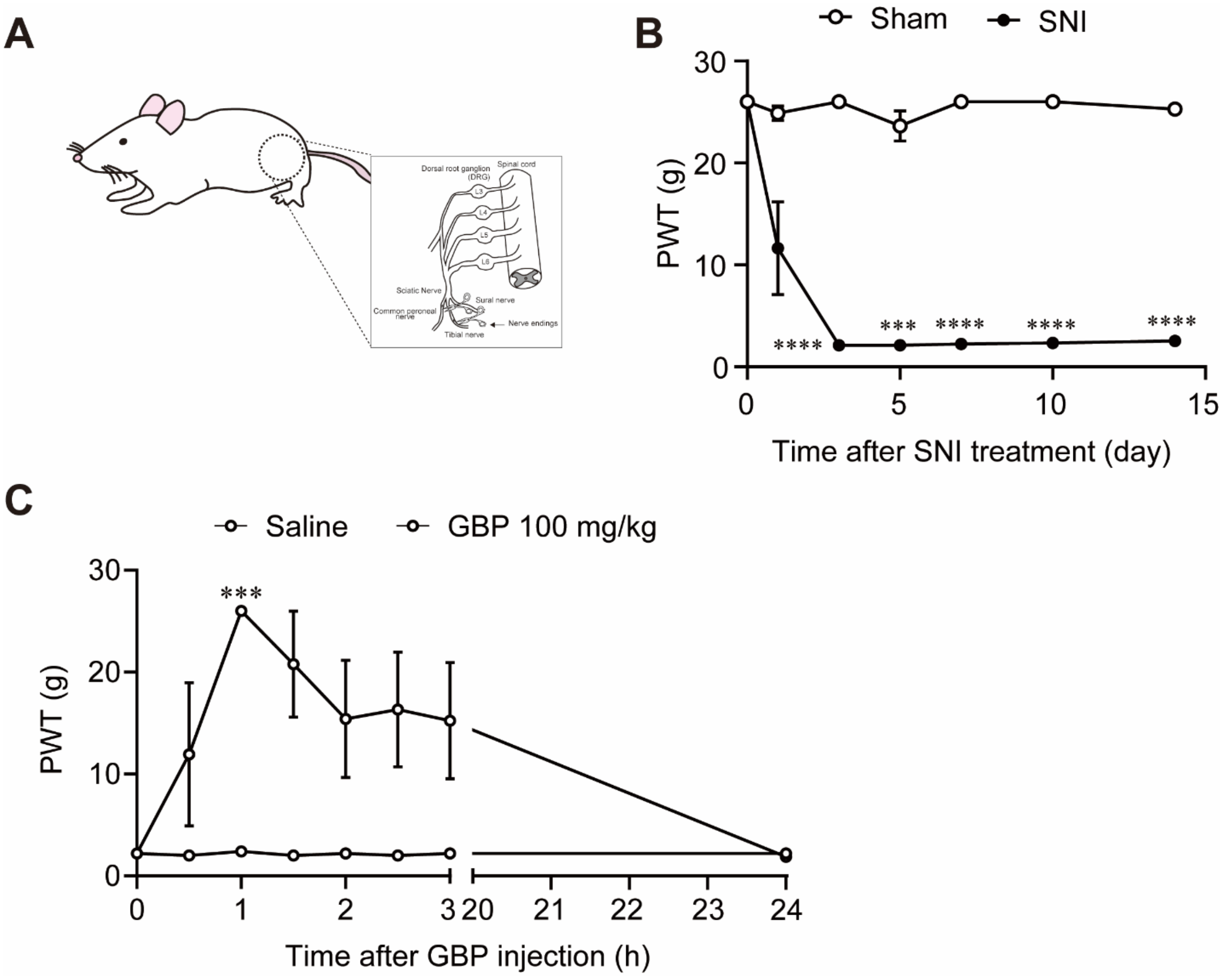


**Fig. S5. Characterization of SNI neuropathic pain rat model.**
(**A**) Configuration of SNI (**B**) long-lasting pain hypersensitivity in SNI treatment. (**C**) Time-dependent changes of anti-hypersensitivity effect in GBP (100 mg/kg, i.p.) injection. ****P*<0.001, *****P*<0.0001. All statistical information details were presented in supplementary table.

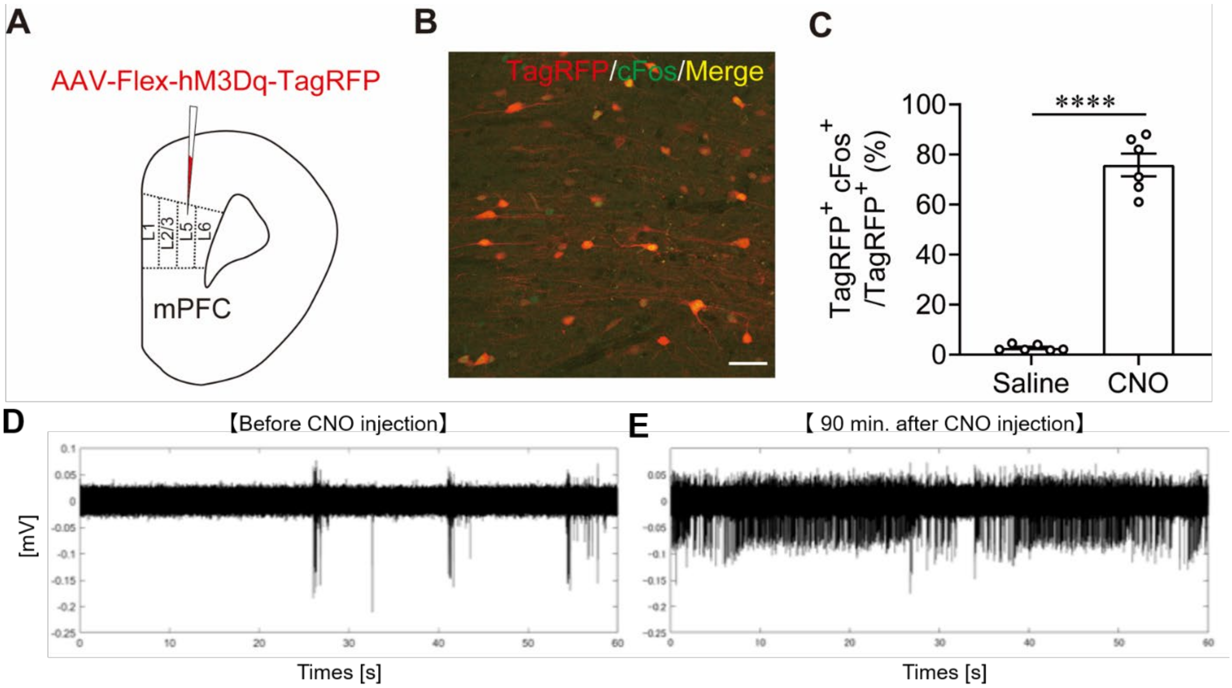


**Fig. S6. Evidence of neural activation by CNO injection in a Gq-DREADD experiment.** (**A**) Experimental procedure for AAV-Flex-hM3Dq-TagRFP injection (**B**) Immunostaining image of TagRFP (red) and cFos (green) at 90 min after saline injection in MOR-Cre KI rat injected AAV-Flex-hM3Dq-TagRFP into mPFC. Scale bar: 50 µm. (**C**) Immunostaining image of TagRFP (red) and cFos (green) at 90 min after Clozapine-*N*-oxide (CNO; 1 mg/kg, i.p.) injection in MOR-Cre KI rat injected AAV-Flex-hM3Dq-TagRFP into mPFC (**D**) Ratio (%) of co-express with TagRFP$^{+}$ neuron (MOR$^{+}$ neuron) and cFos.(**E**) Single unit activities of MOR-Cre KI rat in mPFC before CNO injection (**F**) Single unit activities of MOR-Cre KI rat in mPFC 90 min after CNO (1 mg/kg, i.p.) injection. *****P*<0.0001. All statistical information details were presented in supplementary table.

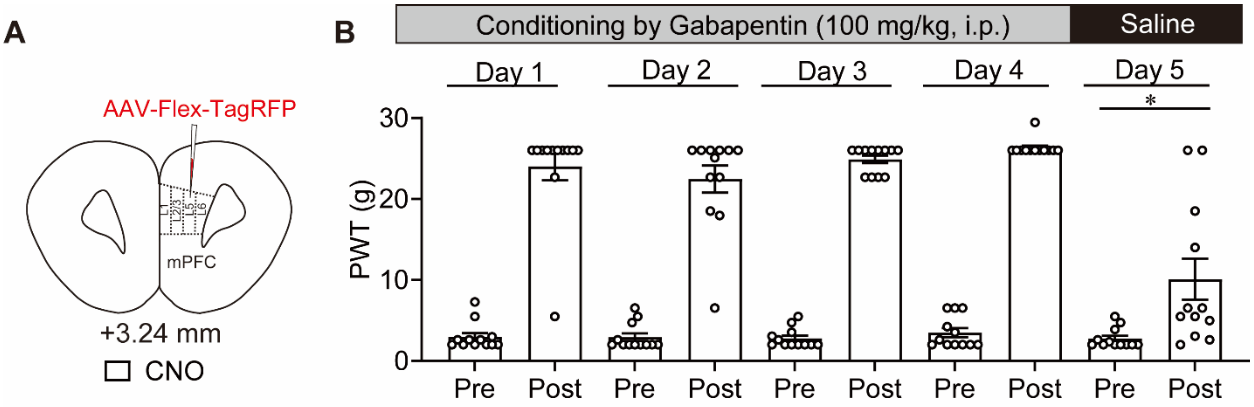


**Fig. S7. Control experiment for Gq-DREADD in conditioning-induced placebo analgesia.** (**A**) Experimental procedure of AAV-Flex-TagRFP (control vector) into mPFC for placebo experiments. (**B**) Change of PWT at 60 min after GBP (conditioning, Day 1–Day 4) and saline (placebo test, Day 5). **P*<0.05. All statistical information details were presented in supplementary table.

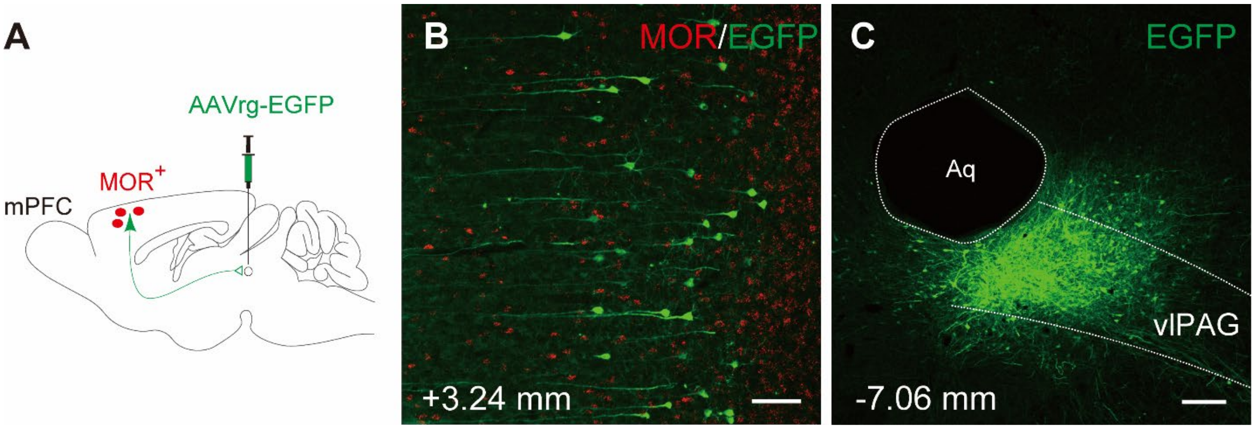


**Fig. S8. Absence of co-expression of MOR and EGFP in L5PN to vlPAG.** (**A**) Experimental procedure for investigating a colocalization with $MOR^+$ neuron (red color) and pyramidal neuron projection vlPAG (showing green color) (**B**) Fluorescent image using RNAscope® and immunohistochemistry of GFP (**C**) Injection area of AAVrg-EGFP in vlPAG. Scale bar: (**B**) 100 µm (**C**) 200 µm.

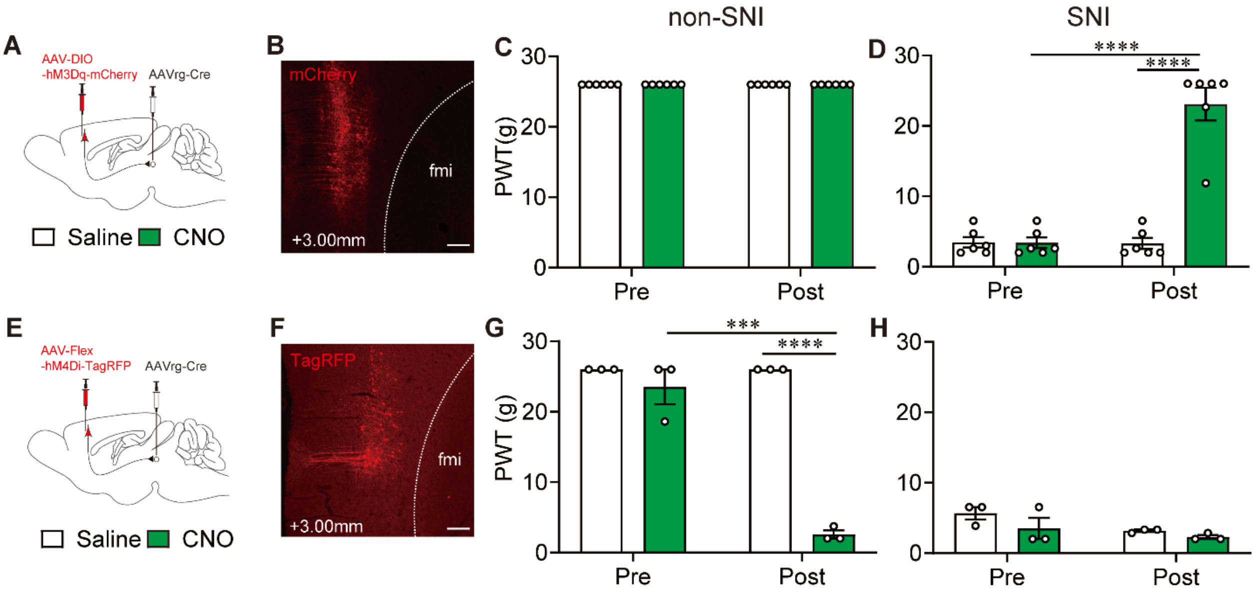


**Fig. S9. Involvement of mPFC-vlPAG pathway in pain modulation.** (**A**) Experimental procedure for Gq-DREADD in non-SNI and SNI rats. (**B**) Immunostaining image for expression of mCherry in mPFC. (**C and D**) Change of PWT after Saline (white column) and CNO, 1 mg/kg, i.p. (green column) in non-SNI (**C**) and SNI (**D**) rats injected AAV-DIO-hM3Dq-mCherry. (**E**) Experimental procedure for Gi-DREADD in non-SNI and SNI rats. (**F**) Immunostaining image for expression of TagRFP in mPFC. (**G and H**) Change of PWT after Saline (white column) and CNO, 1 mg/kg, i.p. (green column) in non-SNI (**G**) and SNI (**H**) rats with AAV-Flex-hM4Di-TagRFP. Scale bar: 200 µm. ***$P$<0.001, ****$P$<0.0001. All statistical information details were presented in supplementary table.

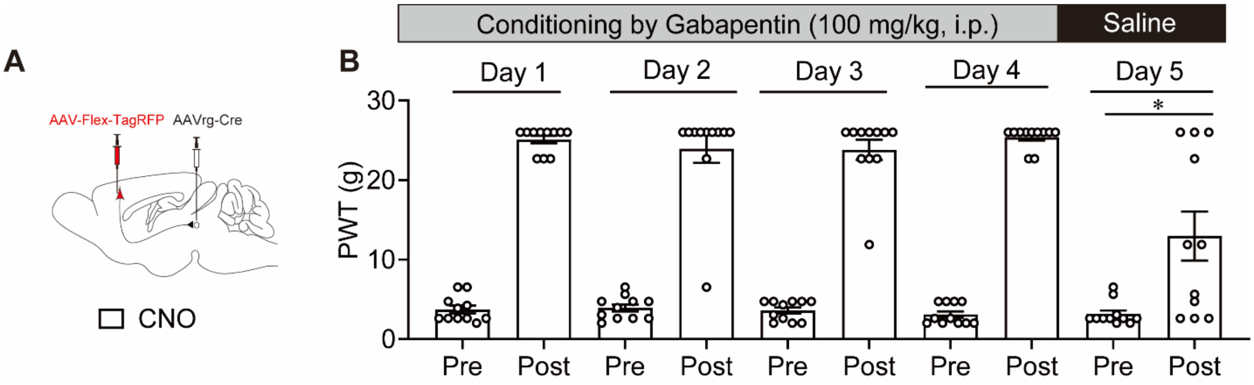


**Fig. S10. Control experiment for Gi-DREADD in conditioning-induced placebo analgesia.** (**A**) Experimental procedure of AAV-Flex-TagRFP (control AAV vector) into mPFC for placebo experiments. (**B**) Change of PWT at 60 min after GBP (conditioning, Day 1–Day 4) and saline (placebo test, Day 5). **P*<0.05. All statistical information details were presented in supplementary table.

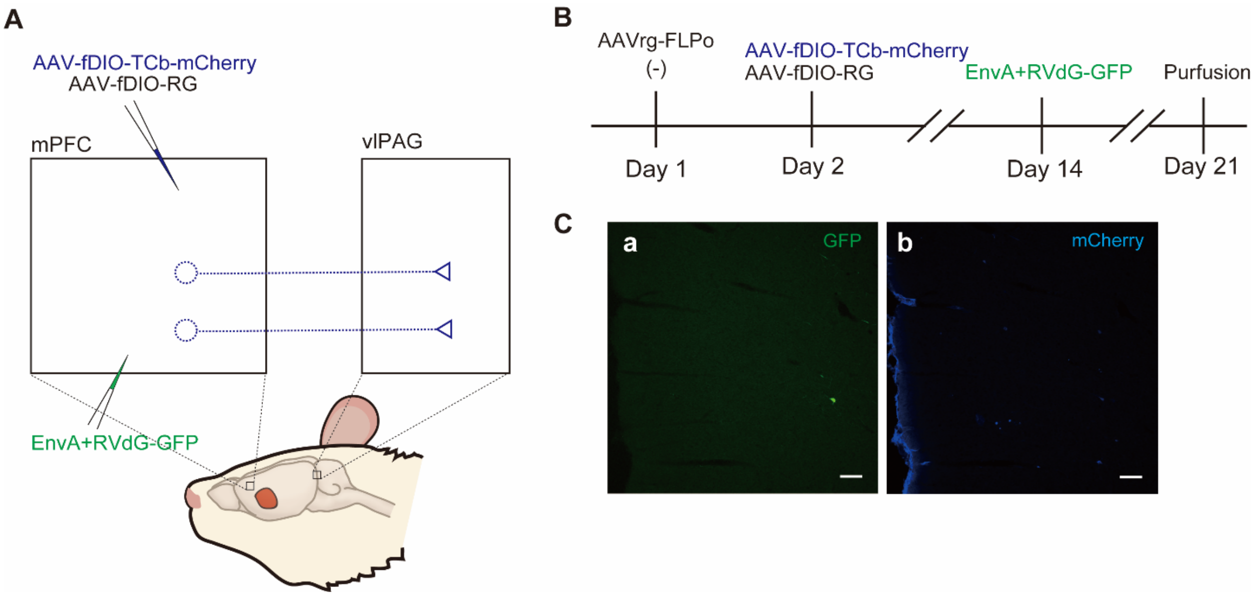


**Fig. S11. Control experiment for TRIO in Fig. 3.** (**A**) Experimental procedures in control experiment of TRIO (**B**) Experimental schedule for control experiments of TRIO (**C**) Immunostaining image, (**a**) GFP (**b**) mCherry. Scale bar: 100 µm.

**Table S1. Number of embryos transferred and KI pups obtained.**

| 1-cell embryo electroporated | 2-cell embryo transferred | Pups delivered | KI pups |
|---|---|---|---|
| 100 | 72 | 3 | 1 |

**Table S2. Sequences and sources of primer for genotyping in MOR-Cre KI rat.**

| Primer sequence | Source | Purpose | Expected product size used with |
|---|---|---|---|
| F1: 5'-TGATGGCCCTCATTCTGAGC-3' | Thermo Fisher | KI check sequencing | WT: -, KI: 980 bp, *R1 |
| F2: 5'-GGCGCTAAGGATGACTCTGG-3' | Thermo Fisher | KI check sequencing | WT: - , KI: 398 bp, *R2<br>WT: - , KI: 1145 bp, *R3 |
| F3: 5'-TAGTTGCCAGCCATCTGTTG-3' | Thermo Fisher | KI check sequencing | N/A |
| R1: 5'-TAACTACCTGTTTTGCCG GG-3' | Thermo Fisher | KI check sequencing | WT: -, KI: 980 bp, *F1 |
| R2: 5'-GACAGCAAGGGGGAGGAT-3' | Thermo Fisher | KI check sequencing | WT: - , KI: 398 bp, *F2 |
| R3: 5'-ATGTGGGGGTTAGGAGAGGG -3' | Thermo Fisher | KI check sequencing | WT: - , KI: 1145 bp, *F2<br>WT: - , KI: 565 bp, *F3 |

**Table S3. Resource and titer of all virus used this study.**

| Name | Titer | Resource | Catalog Number | Figure |
|---|---|---|---|---|
| AAV2-EF1α-Flex-hM3Dq-2A-cgfTagRFP | $2.77\times10^{12}$ copies/ml | Fukushima medial Univ. Dr. Kobayashi Lab. | None | 1, S6 |
| AAV2-EF1α-Flex-hM4Di-2A-cgfTagRFP | $2.7\times10^{12}$ copies/ml | Fukushima medial Univ. Dr. Kobayashi Lab. | None | 1, 2, 5, S9 |
| AAV2-EF1α-Flex-2A-cgfTagRFP | $1.02\times10^{13}$ copies/ml | Fukushima medial Univ. Dr. Kobayashi Lab. | None | S4, S7, S10 |
| AAVrg-CAGGS-IL-2α-EGFP | $2.59\times10^{12}$ copies/ml | Fukushima medial Univ. Dr. Kobayashi Lab. | None | 5 |
| AAV2-EF1α-Flex-ChR2(H134R)-mCherry (36) | $7.98\times10^{12}$ copies/ml | Fukushima medial Univ. Dr. Kobayashi Lab. | None | 4 |
| AAVrg-EF1α-FLPo | $1.1\times10^{13}$ gp/ml | Riken BDR Dr. Miyamichi Lab. | None | 3 |
| AAV5-CAG-FLEx (FRT)-TC-mCherry | $2.6\times10^{12}$ gp/ml | Addgene (MA, USA) | #67827 | 3, S11 |
| AAV8-CAG-FLEx (FRT)-G | $1.3\times10^{12}$ gp/ml | Addgene (MA, USA) | #67828 | 3, S11 |
| Rabies dG-GFP+EnvA | $1.0\times10^{9}$ ip/ml | Riken BDR Dr. Miyamichi Lab | None | 3, S11 |
| AAV5-CaMKIIα-EGFP | $\geqq 3.0\times10^{12}$ ip/ml | Addgene (MA, USA) | #50469 | 2 |
| AAV2-hSyn-DIO-hM3Dq-mCherry | $\geqq 6.0\times10^{12}$ ip/ml | Addgene (MA, USA) | #44361 | S9 |
| AAVrg-pgk-Cre | $\geqq 7.0\times10^{12}$ ip/ml | Addgene (MA, USA) | #24593 | 2, S10 |
| AAVrg-hSyn-EGFP | $\geqq 7.0\times10^{12}$ ip/ml | Addgene (MA, USA) | #50465 | S8 |

gp: genome particle, ip: infectious particles, vg: vector genome

**Table S4. Statistical information for all figures.**

| Figure Number | | Primary | Post-hok | Number of animals |
|---|---|---|---|---|
| Fig. 1F | SNI (-) Stimuli (+)<br>*vs.*<br>SNI (+) stimuli (+)<br>SNI (+) Stimuli (-)<br>*vs.*<br>SNI (+) stimuli (+) | One-way ANOVA | Tukey | SNI (-) Stimuli (+)<br>n=3<br>SNI (+) Stimuli (-)<br>n=5<br>SNI (+) stimuli (+)<br>n=6 |
| Fig. 1H | Post group<br>Saline *vs.* CNO<br>CNO group:<br>Pre *vs.* Post | Two-way RM ANOVA | Bonferroni | Saline n=8<br>CNO n=7 |
| Fig. 1K | Post group<br>Saline *vs.* CNO<br>CNO group<br>Pre *vs.* Post | Two-way RM ANOVA | Bonferroni | Saline n=6<br>CNO n=6 |
| Fig. 1N | Post group<br>Saline *vs.* CNO<br>Saline group<br>Pre *vs.* Post | Two-way RM ANOVA | Bonferroni | Saline n=10<br>CNO n=11 |
| Fig. 2M | Post group<br>Saline *vs.* CNO<br>Saline group<br>Pre *vs.* Post | Two-way RM ANOVA | Bonferroni | Saline n=12<br>CNO n=12 |
| Fig. 5H | PBS *vs.* ITX | Unpaired t-test | | PBS n=5<br>ITX n=6 |
| Fig. 5K | Post group<br>PBS/hM4Di<br>*vs.*<br>ITX/hM4Di<br>PBS/hM4Di group<br>Pre *vs.* Post | Two-way RM ANOVA | Bonferroni | PBS/hM4Di n=5<br>ITX/hM4Di n=6 |
| Fig. S5B | Day 3: Sham *vs.* SNI<br>Day 7: Sham *vs.* SNI<br>Day 10: Sham *vs.* SNI<br>Day 14: Sham *vs.* SNI | Two-way RM ANOVA | Bonferroni | Sham n=5<br>SNI n=5 |
| Fig. S5C | 1 hour after injection<br>Saline<br>*vs.*<br>GBP 100 mg/kg | Two-way RM ANOVA | Bonferroni | Saline n=3<br>GBP n=3 |
| Fig. S6C | Saline *vs.* CNO | Unpaired t test with Welch's correction | | Saline n=6<br>CNO n=6 |
| Fig. S7B | Day 5: Pre *vs.* Post | Unpaired t test with Welch's correction | | n=12 |
| Fig. S9D | Post group<br>Saline *vs.* CNO<br>CNO group<br>Pre *vs.* Post | Two-way RM ANOVA | Bonferroni | Saline n=6<br>CNO n=6 |
| Fig. S9G | Post group<br>Saline *vs.* CNO<br>CNO group<br>Pre *vs.* Post | Two-way RM ANOVA | Bonferroni | Saline n=3<br>CNO n=3 |
| Fig. S10B | Day 5<br>Pre *vs.* Post | Unpaired t test with Welch's correction | | n=11 |
